# Cancer cell phagocytosis induces an anti-inflammatory gene regulatory program in macrophages

**DOI:** 10.1101/2025.08.18.670953

**Authors:** Catherine R. Zhang, Lei Xiong, Mingxin Gu, Christine Y. Yeh, Livnat Jerby, Anshul Kundaje, William J. Greenleaf, Michael C. Bassik

**Affiliations:** Cancer Biology Program, Stanford University, Stanford, CA, US; Department of Genetics, Stanford University, Stanford, CA, US; Department of Applied Physics, Stanford University, Stanford, CA, US; Department of Computer Science, Stanford University, Stanford, CA, US

## Abstract

Macrophages are capable of eliminating cancer cells by phagocytosis, particularly in the presence of monoclonal antibody (mAb) therapies targeting tumor antigens. Paradoxically, tumor-associated macrophages are typically associated with poor patient outcome, and can promote tumor growth by secretion of immunosuppressive cytokines and growth factors. The mechanisms by which these pro-tumor macrophage states arise are poorly understood, and it is unclear how mAb-induced cancer cell phagocytosis may contribute to these states. To understand how antibody-dependent cancer cell phagocytosis (ADCP) alters macrophage state and function, we profiled gene expression and chromatin accessibility changes over time after ADCP. We observed that after ADCP, macrophages upregulate an anti-inflammatory gene regulatory program, characterized by expression of pro-angiogenic and immunosuppressive chemokine genes, and increased activity by cellular, oxidative, and lysosomal stress transcription factors. This gene regulatory program was shared among phagocytic macrophages following either ADCP or apoptotic cancer cell phagocytosis, in addition to substrate-specific pathways. Conditioned media from macrophages promoted EMT in cancer cells, but this pro-EMT macrophage phenotype was attenuated following ADCP, but not following apoptotic cancer cell phagocytosis. The phagocytic gene signature we identified *in vitro* is also expressed by tumor-associated macrophages across numerous cancer types *in vivo*. Together, this work identifies an anti-inflammatory and immunosuppressive epigenetic program in macrophages following ADCP upon mAb treatment, and expands our understanding of how phagocytosis influences macrophage heterogeneity in the tumor microenvironment.

## Introduction

Macrophages are highly plastic myeloid cells that play key roles in inflammation, wound healing, and disease. Across these diverse contexts, macrophages detect and phagocytose an array of extracellular substrates, including pathogens and apoptotic cells, and phagocytosis of these different substrates can induce either inflammatory or immunosuppressive responses in macrophages.^1–3^ For example, Fcγ receptor-mediated phagocytosis of IgG antibody-opsonized pathogens activates ITAM-Syk signaling pathways in macrophages that trigger transcription of pro-inflammatory genes.^3–7^ In contrast, during wound healing, efferocytosis, or phagocytosis of apoptotic cells, induces an anti-inflammatory and reparative gene expression program in macrophages.^8–12^

In the context of cancer, macrophages are often the most abundant immune cell type in the tumor microenvironment (TME), yet their infiltration is typically associated with poor prognosis in cancer.^13–17^ Although macrophages are capable of mounting anti-tumor responses through phagocytosis of cancer cells and antigen presentation to adaptive immune cells, most tumor-associated macrophages (TAMs) occupy a pro-tumor, anti-inflammatory state, where they secrete immunosuppressive cytokines and growth factors, promote angiogenesis, and help establish a pre-metastatic niche.^18–21^ Macrophages and dendritic cells also phagocytose apoptotic cancer cells and cancer cell debris, which induces expression of immunosuppressive genes.^22–24^ However, while numerous studies have profiled the transcriptional and epigenetic heterogeneity of TAMs *in vivo,*^25–31^ the mechanisms underlying the observed pro-tumor states of macrophages, particularly their modulation by phagocytosis, remain incompletely understood.

Understanding the functional impacts of cancer cell phagocytosis on macrophage state is further complicated by the growing application of therapeutic monoclonal antibodies (mAbs) (e.g. anti-CD20, anti-HER2, and anti-CD47), which bind the cancer cell antigen, induce subsequent recognition of the mAb by the Fcγ receptor (FcγR) on macrophages, and lead to antibody-dependent cellular phagocytosis (ADCP).^32–37^ Although mAb treatment induces cancer cell killing by macrophages, mAb therapy still faces challenges in non-response, durability, and resistance, perhaps as a consequence of ADCP itself. Consistent with this hypothesis, ADCP macrophages cultured with breast cancer cell lines in the presence of Trastuzumab (anti-HER2) upregulate PD-L1 surface expression, inhibiting NK-and T-cell mediated cytotoxicity,^38^ suggesting that ADCP macrophages may exhibit an immunosuppressive effect in the TME. Therefore, while macrophages in the TME are capable of eliminating both apoptotic and live cancer cells by phagocytosis, particularly in the context of mAb therapies, it is not well understood how these distinct types of phagocytosis shape macrophage molecular state and ultimately their effect on tumors. A detailed molecular understanding of how ADCP alters macrophage state could elucidate mechanisms of non-response and resistance to mAb therapy.

Here, we investigate the link between mAb-associated ADCP and pro-tumor macrophage states, decoupling the effects of distinct cell phagocytosis substrates and identifying molecular mechanisms by which cancer cell phagocytosis alters macrophage state. To this end, we systematically profile gene expression and chromatin accessibility changes in macrophages over time following ADCP of mAb-opsonized live cancer cells *in vitro* and compare these changes to those induced by efferocytosis. Our study demonstrates that although mAb treatment leads to cancer cell phagocytosis and induces expression of antigen presentation genes in macrophages, ADCP also induces expression of immunosuppressive genes including *Cd274* (PD-L1), *Gpnmb, Spp1*, and *Trem2* as well as pro-angiogenic genes including *Vegfb*, *Angpt2*, and *Mmp9*. By comparing the effect of ADCP and efferocytosis in the same *in vitro* system, we identify a conserved cellular, lysosomal, and oxidative stress TF response program induced by both forms of phagocytosis, as well as substrate-specific pathways. In addition to the effect of ADCP on macrophage molecular state, we demonstrate that ADCP uniquely alters the secretome of macrophages, and conditioned media from ADCP macrophages leads to upregulation of immunosuppressive chemokines in cancer cells and lower expression of EMT genes compared to macrophage conditioned media. Finally, we provide evidence that these cancer cell phagocytosis gene signatures occur in TAMs across numerous cancer types *in vivo*.

## Results

### ADCP induces an anti-inflammatory and immunosuppressive gene expression program in macrophages

To capture the transcriptional and epigenetic changes in macrophages following ADCP of live cancer cells, we developed an approach to sort macrophages in a co-culture system based on whether they had undergone ADCP, and then profiled the sorted populations by RNA- and ATAC-seq (Fig. 1a). To distinguish macrophages that had phagocytosed a live cancer cell from those that had not, we labeled human Ramos lymphoma cells with both calcein, a live cell stain, and pHrodo red, a pH-sensitive fluorophore that increases in signal after the cell reaches the acidic pH of the macrophage lysosome. We then opsonized these labeled Ramos cells with anti-CD20 and anti-CD47 mAbs to induce ADCP. To enable us to distinguish whether transcripts originated from phagocytosed cancer cells or macrophages, we co-cultured pHrodo- and calcein-labeled human Ramos cells with mouse macrophages. We selected the mouse macrophage cell line J774 because these cells efficiently phagocytose live cancer cells in the presence of anti-CD20 and anti-CD47, and have previously been used as a model to study live cancer cell phagocytosis.^39^ During analysis, we then discarded sequencing reads that mapped to the human genome to retain only transcripts from mouse macrophages.

**Figure 1.**
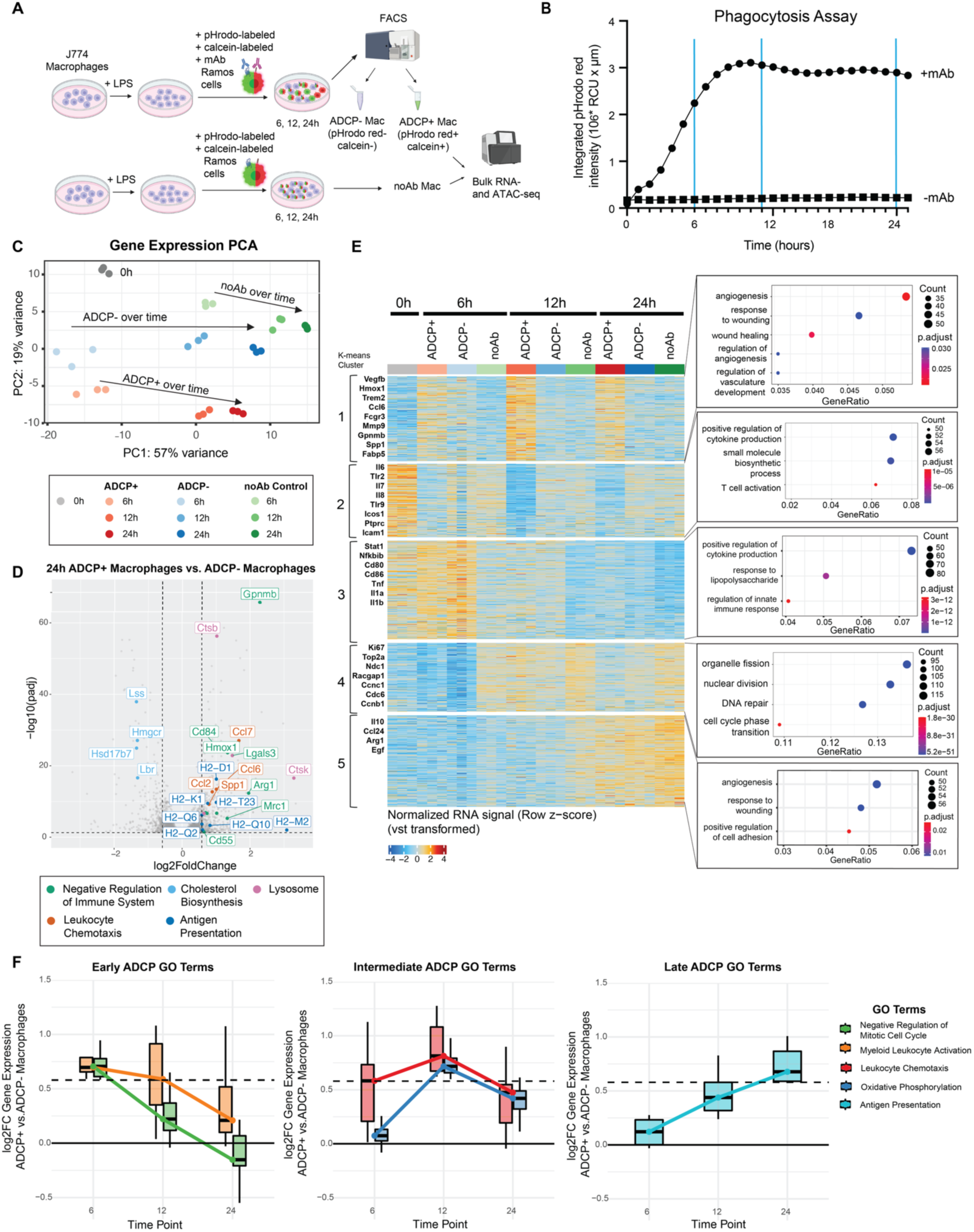
Antibody-dependent cellular phagocytosis (ADCP) induces an anti-inflammatory and immunosuppressive gene expression program in macrophages. a. Schematic of time course sorting J774 macrophages based on ADCP of live cancer cells, and performing RNA- and ATAC-seq (n = 3 biological replicates per condition). b. Phagocytosis assay quantifying pHrodo red signal (which increases upon acidification in the lysosome) over time after co-culture with Ramos cells +/- mAb with selected time points for time course. c. Principal component analysis (PCA) of gene expression profiles of ADCP+, ADCP-, and noAb macrophages at each time point. d. Differentially expressed genes (|log2FC| > 0.58, p-adj < 0.05) between ADCP+ and ADCP-macrophages at 24 hours. Color of labeled points corresponds to GO term. e. K-means clustering of ADCP+, ADCP-, and no Ab control J774 macrophages at 6, 12, and 24 hours, visualized as a heatmap of gene expression. Each row represents a z-score of vst-transformed counts of gene expression. GO term enrichment on genes in each cluster over all differentially expressed genes from k-means clustering. f. Enriched GO terms at early, intermediate, and late time points following ADCP. Each boxplot represents differential expression of genes in each enriched GO term between ADCP+ and ADCP-macrophages at each time point.

To identify time-dependent gene regulatory changes induced by ADCP, we co-cultured lipopolysaccharide (LPS)-activated macrophages with Ramos cells in the presence or absence of mAb for 6, 12, and 24 hours. At each time point, we sorted macrophages co-cultured with cancer cells in the presence of mAb based on their calcein and pHrodo signal into ADCP+ macrophages (pHrodo+ calcein+) and ADCP-macrophages (pHrodo-calcein-) (Fig. 1a; Fig. S1a). We also collected macrophages that were co-cultured with Ramos cells in the absence of mAb (noAb) to control for cancer cell secreted factors. Phagocytosis of live cancer cells peaked at 12 hours, plateaued by 24 hours, and was dependent on the presence of mAb (Fig. 1b). Sorted macrophages were then lysed for RNA extraction and RNA-seq (Fig. S1b-c) or prepared for ATAC-seq.

While PCA revealed that the primary source of variance (57% of variance) in gene expression profiles across conditions was driven by co-culture time with cancer cells and washing out of the initial LPS stimulus, PC2 (19% of variance) clearly separated cells based on whether or not they had undergone ADCP (Fig. 1c). To determine how ADCP alters gene expression in macrophages, we first examined differentially expressed genes between ADCP+ and ADCP-macrophages at the last time point (24 hours) (Fig. 1d; Supplementary Table 1). In ADCP+ macrophages, we observed an upregulation of lysosomal genes such as *Ctss* and *Ctsk*, consistent with previous reports of lysosomal gene upregulation following FcγR activation.^40^ We also observed an upregulation of lipid metabolism genes (*Fabp5*), MHC Class I antigen presentation genes (*H2-K1* and *H2-D1*) as well as immunosuppressive genes (*Gpnmb, Cd274/*PD-L1*, Spp1,* and *Trem2*) and canonical anti-inflammatory markers (*Mrc1/*CD206 and *Arg1*). Thus, while ADCP induces MHC Class I antigen presentation genes, supporting prior reports of CD8+ T cell priming by ADCP macrophages,^41^ ADCP also upregulates immunosuppressive secreted factors and ligands, revealing a dual functional state in macrophages that may underlie their ability to inhibit NK and T cell cytotoxicity.^38^

To identify global patterns of gene expression changes following ADCP, we then combined all differentially expressed genes across all pairwise comparisons and conducted k-means clustering on this gene set (Supplementary Table 2). K-means clustering revealed the effect of ADCP (cluster 1-2) and increasing co-culture time (clusters 3-5) (Fig. 1e; Supplementary Text; Supplementary Table 2). Cluster 1 (*Gpnmb+ Spp1+*), which was composed of genes with higher expression in ADCP+ macrophages at every time point, was enriched for genes associated with phagocytosis such as *Fcgr3* as well as genes associated with wound healing and angiogenesis pathways, including *Hmox1, Vegfb*, and *Mmp9.*^42–44^ Cluster 1 (*Gpnmb*+ *Spp1*+) also included genes linked to immunosuppressive macrophages whose infiltration is associated with poor prognosis in cancer, including *Mrc1 (*CD206)*, Spp1, Gpnmb,* and *Trem2.*^25,45–50^ Cluster 2 (*Il6*+ *Tlr2*+), which contained genes with lower expression in ADCP+ compared to ADCP-macrophages at later time points, was enriched for genes in cytokine production and T cell activation pathways, including *Il6, Il7, Il8*, and *Tlr2*, suggesting that ADCP+ macrophages express lower levels of pro-inflammatory genes. Thus, we observe that ADCP leads to the upregulation of pro-angiogenic and anti-inflammatory genes and downregulation of pro-inflammatory genes in macrophages.

Next, we leveraged our time course to investigate the time-dependent gene expression programs induced by ADCP, comparing differentially expressed genes between ADCP+ and ADCP-macrophages at each time point (Fig. S1c-d). After 6 hours of co-culture, ADCP+ macrophages showed increased expression of genes associated with myeloid leukocyte activation (*Cd33* and *Il10)* and mitotic spindle rearrangement (*Racgap1, Arhgef7, Cdkn2c,* and *Wee1)* (Fig. 1f). Beyond their role in cell cycle, *Racgap1* and *Arhgef7* regulate Rho GTPases, facilitating the actin cytoskeletal remodeling required for phagocytic cup formation.^51–53^ The increased expression of these gene programs in ADCP+ compared to ADCP-macrophages was most pronounced early on and declined over time, aligning with their role in initiating phagocytosis (Fig. 1f).

We observed the greatest number of differentially expressed genes between ADCP+ and ADCP-macrophages after 12 hours of co-culture (Fig. S1c). Genes with higher expression at 12 hours in ADCP+ macrophages were enriched for oxidative phosphorylation genes, including NADH dehydrogenases, ATPase subunits, and cytochrome C oxidase subunits, as well as leukocyte chemotaxis genes, including *Ccl8, Ccl6, Ccl2,* and *Ccl7* (Fig. 1f). The induction of oxidative phosphorylation following ADCP is consistent with previous studies *in vivo* that demonstrated that TAMs that have phagocytosed neoplastic cells shift to oxidative phosphorylation metabolism,^23^ which has also been associated with immunosuppressive phenotypes in macrophages.^54–56^

After 24 hours of co-culture, ADCP+ macrophages expressed higher levels of genes associated with MHC Class I antigen presentation (Fig. 1f). At 24 hours, ADCP+ macrophages also upregulated inhibitors of leukocyte-mediated activation, including inhibitors of the complement membrane attack complex (*Cd55* and *Cd59a*), regulators of T cell exhaustion (*Cd274*/PD-L1), and granzyme B inhibitor (*Serpinb9b*). 25 genes were consistently upregulated in ADCP+ macrophages compared to ADCP-macrophages at every time point (Fig. S1e, Supplementary Table 1), including anti-inflammatory genes *Mrc1, Arg1, Marco, Hmox1,* and *Gpnmb (*Fig. 1f). Collectively, these results suggest that ADCP induces an anti-inflammatory state in macrophages, characterized by upregulation of immunosuppressive chemokine and pro-angiogenic genes and downregulation of pro-inflammatory cytokine genes.

### Cellular, lysosomal, and oxidative stress transcription factors regulate ADCP gene expression programs

Macrophages adopt distinct phenotypes in response to external stimuli by adjusting both their gene expression and epigenetic programs.^57–59^ Since we observed distinct changes in gene expression in macrophages after ADCP, we asked whether concordant changes in chromatin accessibility reflect regulation of these gene programs. We performed bulk ATAC-seq on the same macrophage populations following ADCP (Fig. S2a-d). Similar to the RNA-seq, the dominant source of variation in PCA on chromatin accessibility data was associated with co-culture time with cancer cells and washing out LPS, but PC2 (12% of variance) clearly separated ADCP+ from ACDP-macrophages at all timepoints (Fig. 2a).

**Figure 2.**
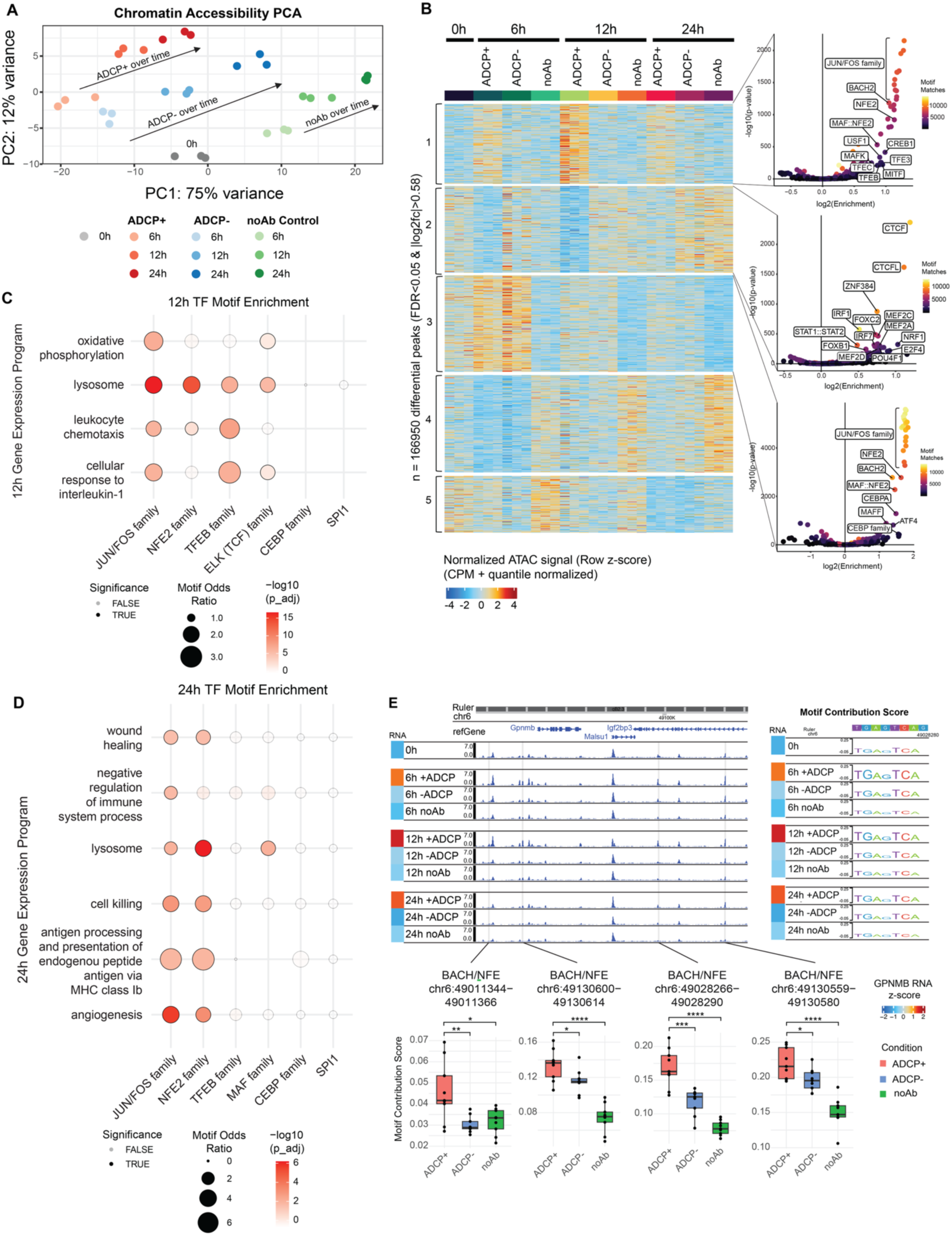
Cellular, lysosomal, and oxidative stress transcription factors regulate ADCP gene expression programs. a. Principal component analysis (PCA) of chromatin accessibility profiles of ADCP+, ADCP-, and noAb macrophages at each time point (n = 3 biological replicates per condition). b. K-means clustering of ADCP+, ADCP-, and no Ab control J774 macrophages at 6, 12, and 24 hours, visualized as a heatmap of chromatin accessibility. Each row represents a z-score of normalized ATAC signal. TF motif enrichment in peaks in each cluster over all differentially accessible peaks. c. Transcription factor family motif enrichment in more accessible peaks in ADCP+ macrophages compared to ADCP-macrophages (log2FC > 1, adjusted p-value < 0.05) within 250kb of genes in GO terms enriched in upregulated genes in ADCP+ macrophages compared to ADCP-macrophages at 12 hours. Motif enrichment was calculated using Fisher’s exact test with unchanging peaks within 250kb of gene programs induced in ADCP+ macrophages as background. d. Transcription factor family motif enrichment in more accessible peaks in ADCP+ macrophages compared to ADCP-macrophages (log2FC > 1, adjusted p-value < 0.05) within 250kb of genes in GO terms enriched in upregulated genes in ADCP+ macrophages compared to ADCP-macrophages at 24 hours. Motif enrichment was calculated using Fisher’s exact test with unchanging peaks within 250kb of gene programs induced in ADCP+ macrophages as background. e. ChromBPNet model-derived contribution scores at BACH/NFE family motifs that are highly correlated (Pearson correlation > 0.5) with *Gpnmb* RNA expression within 100 kb of *Gpnmb* TSS. Statistical significance between contribution scores across ADCP conditions was calculated using a Wilcoxon-rank sum test. Example contribution score of 1 replicate at BACH/NFE family motif is shown on top right. Motif loci are shown on ATAC genome browser tracks on the left.

To examine global patterns of chromatin accessibility changes following ADCP, we combined all differentially accessible elements across all pairwise comparisons and conducted K-means clustering on this peak set. K-means clustering revealed the effect of ADCP (cluster 1-2) and increasing co-culture time (clusters 3-5) (Fig. 2b). To nominate trans-acting transcription factors (TFs) associated with changes in chromatin accessibility, we identified TF motifs enriched in the peaks in each cluster.

Cluster 1 peaks, which were more accessible in ADCP+ macrophages across all time points, were enriched for motifs of cellular, oxidative, and lysosomal stress response transcription factors (TFs) including JUN/FOS, BACH2, NFE2L2 (NRF2), and TFEB. NRF2 regulates oxidative stress,^60–63^ promotes resistance to cell death, and contributes to an immunosuppressive TAM phenotype,^64–66^ which aligns with our finding from transcriptional data that ADCP+ macrophages upregulate expression of immunosuppressive cytokine genes. In response to oxidative stress, NRF2 activates *Hmox1* as part of a cytoprotective response,^67,68^ and we found that *Hmox1* had higher expression in ADCP+ macrophages compared to ADCP-macrophages across time points (Fig. 1f, Supplementary Table 1). Consistent with our observed enrichment of TFEB family motifs in Cluster 1 accessible elements in ADCP+ macrophages, Fcγ-receptor-mediated phagocytosis has been shown to induce TFEB activation and activate lysosomal biogenesis genes in response to lysosomal stress.^40,69^

Cluster 2, composed of peaks with lower accessibility in ADCP+ compared to ADCP-macrophages, was enriched for motifs of CTCF, IRFs, and STAT1/STAT2. IRFs and STAT1 have been shown to activate pro-inflammatory genes upon IFNγ stimulation;^57,70,71^ thus, reduction in activity of these factors is consistent with our transcriptional data showing that ADCP+ macrophages downregulated expression of pro-inflammatory cytokines. Cluster 3, composed of peaks with higher accessibility in macrophages at early time points, was enriched for motifs of JUN/FOS, known regulators of pro-inflammatory genes following LPS stimulation.^72–75^ Notably, while cluster 1 and cluster 3 share an enrichment for JUN/FOS motifs, TFEB family motifs are unique to cluster 1 enrichment. Thus, our analysis suggests that ADCP+ macrophages exhibit increased activity of cellular, oxidative, and lysosomal stress TFs and decreased activity of pro-inflammatory TFs.

Next, we sought to identify transcription factors driving the pro-tumor gene expression programs we observed in ADCP+ macrophages. Using our paired RNA- and ATAC-seq data, we identified nearby putative enhancers whose accessibility was predictive of expression at genes of interest. At each time point, we selected Gene Ontology terms that were enriched in upregulated genes and TF motifs that were enriched in more accessible peaks in ADCP+ macrophages (Fig. S2e; Supplementary Table 1,3). For each induced gene expression program, we identified all nearby peaks within 250 kb of each gene TSS, and quantified the enrichment of each TF motif in nearby more accessible peaks compared to nearby unchanging peaks (Fig. 2c-d). We first quantified the enrichment of macrophage lineage-defining (CEBP family and SPI1) transcription factor motifs, which was not enriched in nearby accessible peaks near ADCP-induced gene expression programs (Fig. 2c-d; Supplementary Table 4).

Because we observed an upregulation of lysosomal genes at 12 hours following ADCP in our RNA data as well as an enrichment for TFEB family motifs in more accessible peaks following ADCP in our ATAC data, we used our peak-gene linkage method to predict the gene expression programs regulated by TFEB, a known activator of lysosome biogenesis genes. As expected, we observed a significant enrichment for TFEB motifs in more accessible peaks near upregulated lysosomal genes (Fig. 2c). Notably, we also observed a significant enrichment of TFEB motifs in more accessible peaks near leukocyte chemotaxis and interleukin-1 response genes (Fig. 2c). At 24 hours, we no longer observed a significant enrichment of TFEB family motifs near induced gene expression programs (Fig. 2d). More accessible peaks at 12 hours in ADCP+ macrophages near lysosomal genes were also enriched for JUN/FOS and BACH/NFE family motifs. JUN/FOS family motifs were enriched in more accessible peaks near all the ADCP-induced gene programs, with the greatest enrichment in differential peaks near oxidative phosphorylation genes. At 24 hours, fewer genes were differentially expressed and fewer peaks were differentially accessible between ADCP+ and ADCP-macrophages compared to at 12 hours (Fig. S1c, S2c). Similar to at 12 hours, JUN/FOS and BACH/NFE family motifs were enriched in nearby more accessible peaks to all induced gene expression programs at 24 hours, with the strongest enrichment near MHC class Ib antigen presentation genes (Fig. 2d; Supplementary Table 4). Collectively, our integrated RNA- and ATAC-seq analysis demonstrates that the upregulated ADCP gene expression programs are primarily regulated by JUN/FOS and BACH/NFE family TFs, consistent with recent reports that NRF2 (NFE2L2) contributes to immunosuppressive TAM phenotypes *in vivo*.^64^ While TFEB family TFs are known to activate lysosome biogenesis genes upon lysosomal stress,^40,76,77^ recent work has identified a potential role in regulating inflammation in macrophages.^78^ Our data suggests that in addition to its known role in lysosomal stress response, TFEB activity also regulates immune effector gene expression programs induced by ADCP.

To more precisely define which TFs were driving chromatin accessibility changes following ADCP, we next used our ATAC-seq data to train convolutional neural network models (ChromBPNet) that predict ATAC-seq read counts and profiles at base-resolution from the underlying DNA sequence in accessible regions.^79–82,83^ ChromBPNet yields base-resolution contribution scores reflecting the extent to which each nucleotide predicts chromatin accessibility. Clustering recurrent high-contribution sequences yields motifs, which can then be mapped back to peak regions to identify TF motif instances that are predictive of chromatin accessibility.^80^

We used our model-derived contribution scores to investigate the sequence basis of chromatin accessibility dynamics of ATAC-seq peaks within 100kb of *Gpnmb* (Fig. 2e). We examined *Gpnmb* because it is highly expressed and strongly upregulated in ADCP+ macrophages compared to ADCP-macrophages across all time points, and in our k-means clustering analysis, it clusters with immunosuppressive genes (Fig. 1f). *Gpnmb* is reported to be highly expressed in M2 macrophages and upregulated upon M2 polarization,^84–86^ lysosomal stress, and TFEB activation.^87,88^ *Gpnmb* expression has also been associated with immunosuppressive macrophages *in vivo* whose infiltration is associated with poor prognosis in cancer.

To nominate TF motifs that could be driving *Gpnmb* upregulation following ADCP, we calculated the correlation between *Gpnmb* RNA expression and our model-derived contribution scores at all motif instances within 100kb of the *Gpnmb* TSS. Of the highly correlated (Pearson correlation > 0.5) motifs, BACH/NFE family motifs had the highest contribution scores across conditions (Fig. S2f; Supplementary Table 5). Contribution scores of four nearby BACH/NFE family motif instances were highly correlated (Pearson correlation > 0.5) with *Gpnmb* RNA expression (Fig. 2e). At each of these motif instances, the contribution score was significantly higher in ADCP+ macrophages compared to ADCP- and noAb control macrophages, suggesting that BACH/NFE TF family binding may drive increased chromatin accessibility at these motifs, which is correlated with increased *Gpnmb* RNA expression. While TFEB family transcription factors have been proposed to regulate *Gpnmb* transcriptionally,^77^ we did not find TFEB family motif instances predicted to regulate chromatin accessibility within 100kb of *Gpnmb* (Supplementary Table 5). Since multiple BACH/NFE family TFs bind a similar motif, we next sought to nominate specific family members that could be driving increased chromatin accessibility. NFE2L1 (NRF1) and NFE2L2 (NRF2) are both activators expressed in J774 macrophages (Fig. S2g), suggesting that these TFs may be binding enhancers near *Gpnmb* to drive increased chromatin accessibility and expression following ADCP.

Collectively, our chromatin accessibility data suggest that cellular, lysosomal, and oxidative stress TFs regulate the lysosomal, oxidative phosphorylation, and anti-inflammatory gene expression programs activated in macrophages upon ADCP.

### Substrate-specific and shared anti-inflammatory gene regulatory programs after ADCP or efferocytosis of cancer cells

Previous studies have demonstrated that during wound healing, efferocytosis (apoptotic cell phagocytosis) leads to the induction of an anti-inflammatory and tissue repair gene expression program in macrophages.^8–12^ In the context of cancer, phagocytosis of apoptotic cell debris leads to the upregulation of immunosuppressive genes such as *Gpnmb, Trem2*, and *Spp1* in macrophages.^8,22^ ADCP and efferocytosis engage different phagocytic receptors,^89,90^ which may lead to different downstream epigenetic and transcriptional effects. However, because we observed that live cancer cell ADCP induces a pro-angiogenic and wound healing gene expression program in macrophages featuring many of these same genes (Cluster 1, Fig. 1e), we hypothesized that ADCP may converge on the same gene regulatory programs as efferocytosis, a central function of macrophages during homeostasis. Therefore, we sought to directly compare the gene regulatory effects of ADCP and efferocytosis in our system.

We treated human Ramos lymphoma cells with staurosporine to induce apoptosis (Fig. 3a, S3a) before co-culture with macrophages. Because efferocytosis is more efficient compared to ADCP (Fig. S3b), when we co-cultured apoptotic cells and macrophages at the same co-culture ratio used to sort ADCP+ and ADCP-macrophages, every macrophage phagocytosed an apoptotic cancer cell (Fig. S3c). Therefore, to sort Effero+ (pHrodo red+) and Effero-(pHrodo red-) macrophages, we titered down the ratio of apoptotic Ramos cells to J774 macrophages to 1:1 (low ratio) (Fig. S3c). We additionally co-cultured the apoptotic Ramos cells to J774 macrophages at a 10:1 ratio (high ratio) to sort “Super-Effero” macrophages (top 10% of pHrodo red+ macrophages) to investigate the effect of higher rates of efferocytosis (Fig. S3c). Notably, while human cancer cell ATAC and RNA reads were detected in ADCP+ macrophages, human cancer cell reads were only detected in Super-Effero macrophages, but not Effero+ or Effero-macrophages (Fig. S3d-e).

**Figure 3.**
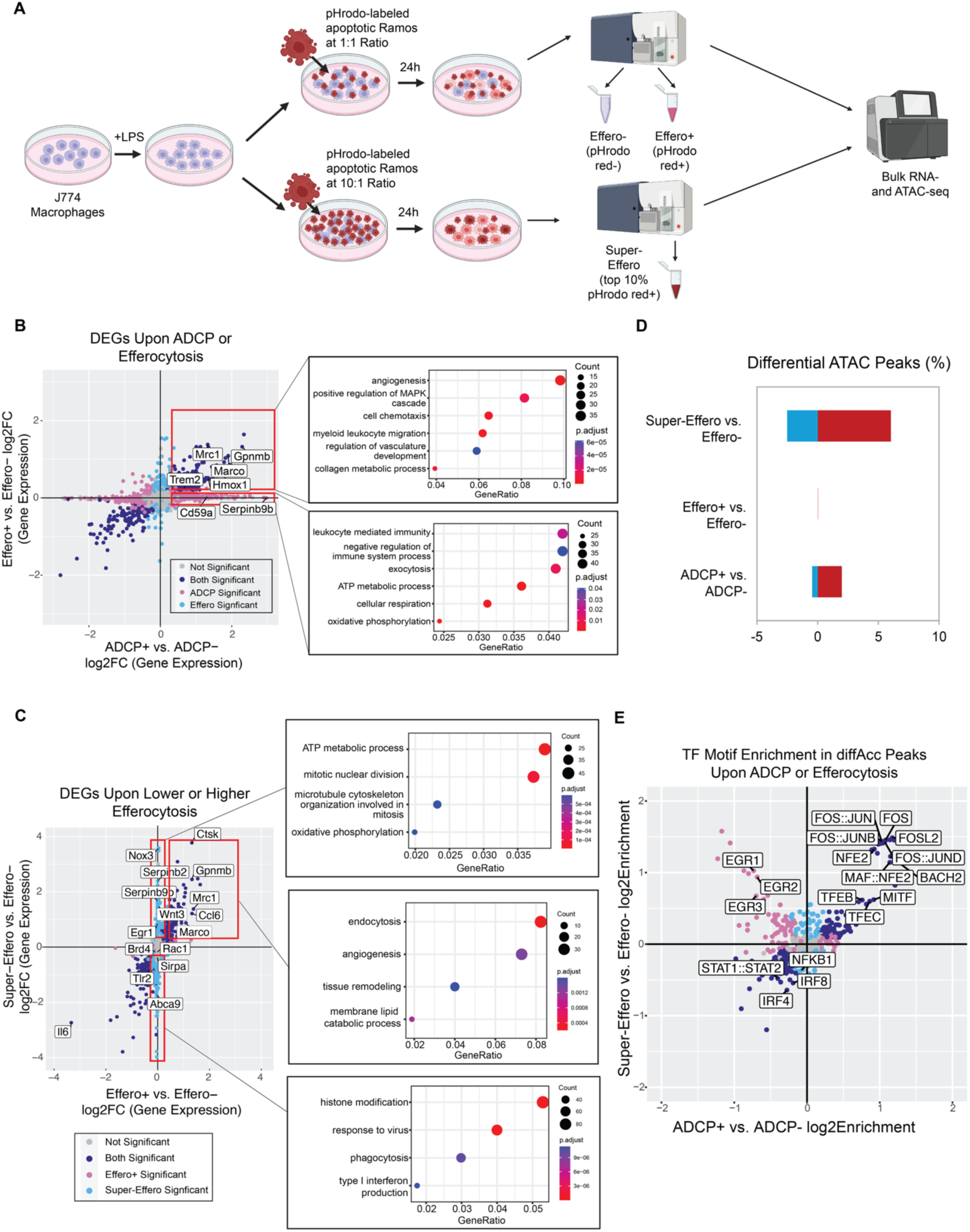
Shared anti-inflammatory gene regulatory program after ADCP or efferocytosis of cancer cells. a. Schematic of co-culturing J774 macrophages at low (1:1) and high (10:1) ratios with apoptotic Ramos lymphoma cells, sorting macrophages based on efferocytosis of apoptotic Ramos lymphoma cells into Effero+, Effero-, and Super-Effero macrophages, and performing bulk RNA- and ATAC-seq. b. Differential gene expression of macrophages upon ADCP (ADCP+ vs. ADCP-log2FC gene expression) or efferocytosis (Effero+ vs. Effero-log2FC gene expression) (n=3 biological replicates per substrate type). Colored points represent significantly differentially expressed genes (p-adj < 0.05). GO term enrichment on genes that are either upregulated in both ADCP+ and Effero+ macrophages, or only upregulated in ADCP+ macrophages. c. Differential gene expression of macrophages upon lower (Effero+ vs. Effero-log2FC gene expression) or higher amounts of efferocytosis (Super-Effero vs. Effero-log2FC gene expression) (n=3 biological replicates per substrate type). Colored points represent significantly differentially expressed genes (p-adj < 0.05). GO term enrichment on genes that are upregulated in both ADCP+ and Super-Effero macrophages, only upregulated in Super-Effero macrophages, or only downregulated in Super-Effero macrophages. d. Percent of ATAC-seq peaks which are differentially accessible (|log2FC| > 0.58, p-adj < 0.05) between ADCP+ vs. ADCP-, Effero+ vs. Effero-, and Super-Effero vs. Effero-macrophages. e. TF motif enrichment in differential accessible peaks (n=3 biological replicates per substrate type) upon ADCP (ADCP+ vs. ADCP-log2Enrichment) or efferocytosis (Effero+ vs. Effero-log2Enrichment). Colored points represent significantly different TF motif enrichment (p-adj < 0.05).

Compared to ADCP+ and ADCP-macrophages, fewer genes were differentially expressed between Effero+ and Effero-macrophages (Fig. S1c, Fig. S3f). Among all genes differentially expressed upon either ADCP or efferocytosis, many genes (17%) were differentially expressed in both conditions with correlated fold changes, but most genes were only differentially expressed upon ADCP (74%) or efferocytosis (9%) respectively (Fig. 3b, Supplementary Table 7). Genes that were upregulated in both ADCP+ and Effero+ macrophages were enriched for genes related to angiogenesis and regulation of vasculature development, including anti-inflammatory macrophage genes and phagocytic receptor genes such as *Mrc1* (CD206) and *Marco* (Fig. 3b). Genes that were upregulated in only ADCP+ macrophages but not Effero+ macrophages were enriched for negative regulators of leukocyte activation, including *Serpinb9b*, inhibitor of granzyme B, and *Cd59a*, an inhibitor of complement activation.

We then compared differential expression of genes upon ADCP or higher levels of efferocytosis exhibited by Super-Effero macrophages. Again, among all genes differentially expressed upon either ADCP or higher levels of efferocytosis, many genes (29%) were differentially expressed in both conditions, but most genes were only differentially expressed upon higher levels of efferocytosis (41%) or ADCP (29%) (Fig. S3g; Supplementary Table 6). Upon higher levels of efferocytosis, *Serpinb9b* was upregulated, suggesting that larger amounts of metabolic stress induced by either higher amounts of efferocytosis or ADCP may lead to a cytoprotective effect in macrophages. *Odc1* (ornithine decarboxylase) was upregulated in Super-Effero macrophages, but unchanged in either Effero+ or ADCP+ macrophages (Fig. S3g; Supplementary Table 6). Consistent with its upregulation only upon higher amounts of efferocytosis, *Odc1* is necessary for continual apoptotic cell internalization, but not apoptotic cell binding, due to its role in metabolism of apoptotic cell-derived arginine and ornithine.^91^ Genes uniquely upregulated in Super-Effero, but unchanged in either Effero+ or ADCP+ macrophages were also enriched for mitotic division genes (Supplementary Table 6). This enrichment is consistent with the proliferative effect of efferocytosis that expands the resolving macrophage population,^10,92^ and suggests that ADCP may not have this same proliferative effect. Thus, we find that macrophages upregulate a shared pro-angiogenic anti-inflammatory gene expression program after ADCP or efferocytosis, in addition to substrate-specific pathways.

During inflammation and injury, macrophages may phagocytose multiple apoptotic cells in succession to maintain homeostasis.^93^ To investigate how the degree of efferocytosis modulates macrophage transcriptional state, we compared differential expression of genes upon lower or higher levels of efferocytosis. Many genes (18%) were correlated in their differential expression upon either lower or higher levels of efferocytosis, with stronger effects observed due to higher efferocytosis. However, most differentially expressed genes (78%) were only differentially expressed upon higher levels of efferocytosis (Fig. 3c; Supplementary Table 6).

Genes that were upregulated in both Effero+ and Super-Effero macrophages were enriched for endocytosis and angiogenesis (Fig. 3c). Genes that were only upregulated in Super-Effero macrophages were enriched for genes in oxidative phosphorylation including NADH dehydrogenase complex subunits, suggesting that metabolic changes in macrophages may only be induced by higher levels of efferocytosis. Genes that were downregulated in only Super-Effero macrophages but not Effero+ macrophages were enriched for genes in phagocytosis, and Type I and II interferon genes (*Ifngr1, Ifnar1, Irf9, Irf7, Tlr4*). While some downregulated genes in Super-Effero macrophages are positive regulators of phagocytosis that are required for actin cytoskeletal rearrangement (*Rac1, Rac2*), Super-Effero macrophages also downregulated *Sirpa* and *Sirpb1*, which are inhibitory receptors that prevent phagocytosis. We also observed an enrichment for chromatin organization (*Ctcf, Brd4, Brd1*) genes in downregulated genes in only Super-Effero macrophages. Consistent with the differential expression of chromatin remodelers only upon higher levels of efferocytosis, in our ATAC-seq data, almost no peaks were differentially accessible between Effero+ and Effero-macrophages, but 8.5% of peaks (16,312 peaks) were differentially accessible between Super-Effero and Effero-macrophages (Fig. 3d).

Next, we compared TF motif enrichment in differentially accessible peaks upon efferocytosis (Super-Effero vs. Effero-macrophages) and ADCP (ADCP+ vs. ADCP-macrophages). Peaks that increased in accessibility in either Super-Effero or ADCP+ macrophages were enriched for motifs of similar transcription factors, including JUN/FOS, TFEB and BACH/NFE family motifs (Fig. 3e; Supplementary Table 7). Peaks that decreased in accessibility in either Super-Effero or ADCP+ macrophages were enriched for motifs of pro-inflammatory TFs such as STAT1/2 and IRFs. Notably, some TF motifs were enriched in peaks that were more accessible upon efferocytosis (Super-Effero vs. Effero-) but less accessible upon ADCP (ADCP+ vs. ADCP-), including EGR family motifs (Fig. 3e; Supplementary Table 7).

Previous studies have demonstrated that EGR1 and EGR3 expression increases rapidly after efferocytosis, but not after FcγR-mediated phagocytosis, to turn on lysosomal acidification and cytoskeleton genes that are necessary for continued efferocytosis.^94^ Consistent with this finding, we observe an increase in *Egr1* expression following higher amounts of efferocytosis, but not ADCP (Supplementary Table 7, Supplementary Fig. 3g).

Collectively, our data suggest that, in addition to substrate-specific pathways, ADCP and efferocytosis induce the upregulation of a similar pro-angiogenic gene expression program, increased activity of shared cellular, lysosomal, and oxidative stress TFs, and decreased activity of shared pro-inflammatory TFs. These results indicate that ADCP unexpectedly induces a reparative and anti-inflammatory gene regulatory program shared with efferocytosis.

### ADCP, but not efferocytosis, attenuates macrophage-induced EMT gene expression programs in cancer spheroids

In the TME, TAMs secrete chemokines and cytokines to recruit and modulate the immune response, and secrete growth factors that promote cancer cell EMT and stemness.^84,95,96^ Previous studies have demonstrated that co-culture with macrophages promotes expression of EMT genes (*Vegfa, Fn1, Itgb1*) and proliferation in lung adenocarcinoma spheroids.^28^ Because both ADCP and efferocytosis induced expression of immunosuppressive chemokines in macrophages, we hypothesized that cancer cell phagocytosis may also alter the macrophage secretome. To test the effect of macrophage secreted factors following ADCP or efferocytosis on cancer cell gene expression, we collected conditioned media (CM) from ADCP+, ADCP-, Effero+, and LPS-stimulated macrophages, and seeded Kras^G12D^ and p53-deficient (KP) mouse lung cancer cell spheroids in methylcellulose-containing CM.^97^ We then conducted RNA-seq on the cancer cell spheroids cultured in different CM (Fig. 4a).

**Figure 4.**
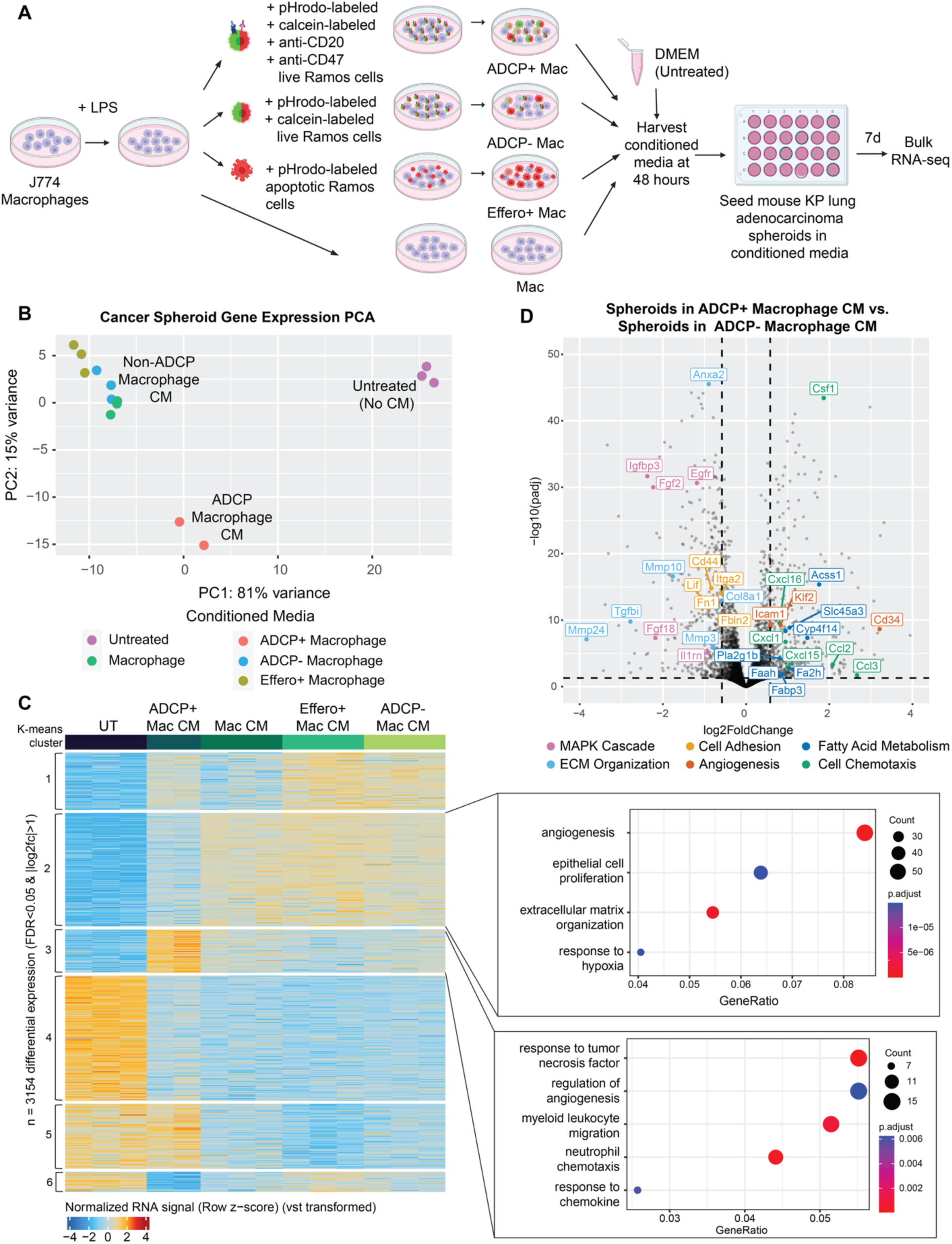
ADCP attenuates macrophage induced-EMT gene expression program in cancer cell spheroids. a. Schematic of mouse lung adenocarcinoma spheroids seeded in conditioned media from macrophages after ADCP, efferocytosis, or live cancer cell co-culture. b. Principal component analysis (PCA) of gene expression profiles of mouse lung adenocarcinoma spheroids seeded in different conditioned media. c. K-means clustering of cancer cell spheroids seeded in conditioned media from ADCP+, ADCP-, Effero+, untreated macrophages, visualized as a heatmap of gene expression. Each row represents a z-score of vst-transformed counts of gene expression. d. Differential gene expression between lung adenocarcinoma spheroids seeded in conditioned media from macrophages co-cultured with mAb-opsonized Ramos cells compared to lung adenocarcinoma spheroids seeded in conditioned media from macrophages co-cultured with Ramos cells. Color of labeled points corresponds to GO term.

Strikingly, conditioned media from macrophages led to large transcriptional changes in lung adenocarcinoma spheroids (Fig. 4b). Spheroids in CM from macrophages, from macrophages that had undergone efferocytosis, or from macrophages that had been co-cultured with live cancer cells (ADCP-macrophages) clustered separately from untreated spheroids (Fig. 4b). However, CM from ADCP+ macrophages induced a transcriptional state distinct from all other macrophage CM-treated spheroids, including Effero+ macrophage CM (Fig. 4b). To identify global patterns of gene expression changes in cancer spheroids in CM from macrophages after ADCP, efferocytosis, or co-culture with cancer cells, we combined all differentially expressed genes across all pairwise comparisons and conducted k-means clustering on this gene set (Fig. 4c). Cluster 2 was composed of genes with low expression in untreated spheroids and high expression in spheroids in macrophage CM, but intermediate expression in spheroids seeded in ADCP+ macrophage CM. Cluster 2 was strongly enriched for genes involved in angiogenesis, epithelial cell proliferation, and ECM remodeling including master regulators of EMT such as *Twist2* and *Sox9*, mesenchymal markers such as *Mmp9, Loxl2,* and *Col8a1*, and growth factors such as *Vegfa, Flt1, Pdgfra,* and *Tgfbi* (Fig. 4c). Cluster 3, which was composed of genes with higher expression in cancer spheroids seeded in ADCP+ macrophage conditioned media, was enriched for genes involved in immune cell chemotaxis such as *Ccl2, Ccl3,* and *Csf1*, as well as genes involved in response to Tnf such as *Tnf* and *Traf1.* These data suggest that ADCP, but not efferocytosis, uniquely alters the macrophage secretome, and that these changes attenuate the pro-EMT molecular phenotype induced by macrophages.

We next focused on the effects of ADCP+ compared to ADCP-macrophages on cancer cell spheroids. We directly compared the gene expression profiles of lung cancer spheroids in CM from macrophages that had been co-cultured with mAb-opsonized live cancer cells (ADCP+ macrophages) to spheroids in CM from macrophages that had only been co-cultured with live cancer cells, but without antibody and thus did not undergo ADCP (ADCP-macrophages) (Fig. 4d; Supplementary Table 8). In spheroids seeded in conditioned media from ADCP+ macrophages, we observed an upregulation of immune chemotaxis genes including *Ccl2* and *Cxcl5*, which recruit immunosuppressive MDSCs,^98,99^ as well as *Csf1*, which induces monocyte-to-macrophage differentiation (Fig. 4d; Supplementary Table 8). Our data suggest a positive feedback loop in which ADCP not only induces the upregulation of immunosuppressive genes in macrophages, but also induces the upregulation of chemokines in cancer spheroids to recruit immune suppressor cells including macrophages. We also observed an upregulation of fatty acid uptake genes such as *Cd36* and *Fabp4* and fatty acid metabolism genes such as *Acss1* and *Slc5a3*. After macrophages take up lipids from phagocytosed cells, they can generate free fatty acids that can then be taken up by other cells in the TME,^100^ including cancer cells, consistent with our observation that cancer cells cultured in ADCP macrophage CM upregulate fatty acid uptake genes. Collectively, our data suggest that macrophage CM induces a pro-EMT gene expression program in cancer cell spheroids. ADCP alters the macrophage secretome to attenuate this pro-EMT gene expression program and leads to the upregulation of immunosuppressive chemokines by cancer cells.

### In vitro ADCP gene signature is expressed by tumor-specific macrophage subset in vivo in mouse models and human cancers

We next asked if the ADCP gene signature we observe *in vitro* was expressed by tumor-associated macrophages in model systems or patient tumors *in vivo*. We first used an existing single-cell RNA-seq dataset of KP mouse models, in which monocytes, interstitial macrophages, and alveolar macrophages were sorted based on phagocytosis *in vivo* of TdTomato+ lung adenocarcinoma cells^23^ (Fig. 5a). Tumor-infiltrating TdTomato+ alveolar macrophages clustered apart from other alveolar macrophage subsets (Fig. 5b). We used our RNA-seq data from our co-culture system to curate a gene module of 155 genes that were upregulated at 24 hours in ADCP+ macrophages compared to ADCP-macrophages (log2FC > 1, adjusted p-value < 0.05) (Supplementary Table 1). We calculated the ADCP gene module score in the alveolar macrophage compartment, which was the predominant phagocytic myeloid cell type *in vivo*^23^ (Fig. 5a). We observed that our ADCP gene signature was significantly more highly expressed in tumor-infiltrating TdTomato+ alveolar macrophages compared to TdTomato-alveolar macrophages (Fig. 5c-d), suggesting that the transcriptomic changes we observed upon ADCP *in vitro* are also induced by cancer cell phagocytosis *in vivo*. Our ADCP gene signature was also significantly more highly expressed in TdTomato+ compared to TdTomato-interstitial macrophages, but not differentially expressed between TdTomato+ and TdTomato-monocytes, suggesting that our ADCP gene signature is macrophage-specific (Fig. S4a).

**Figure 5.**
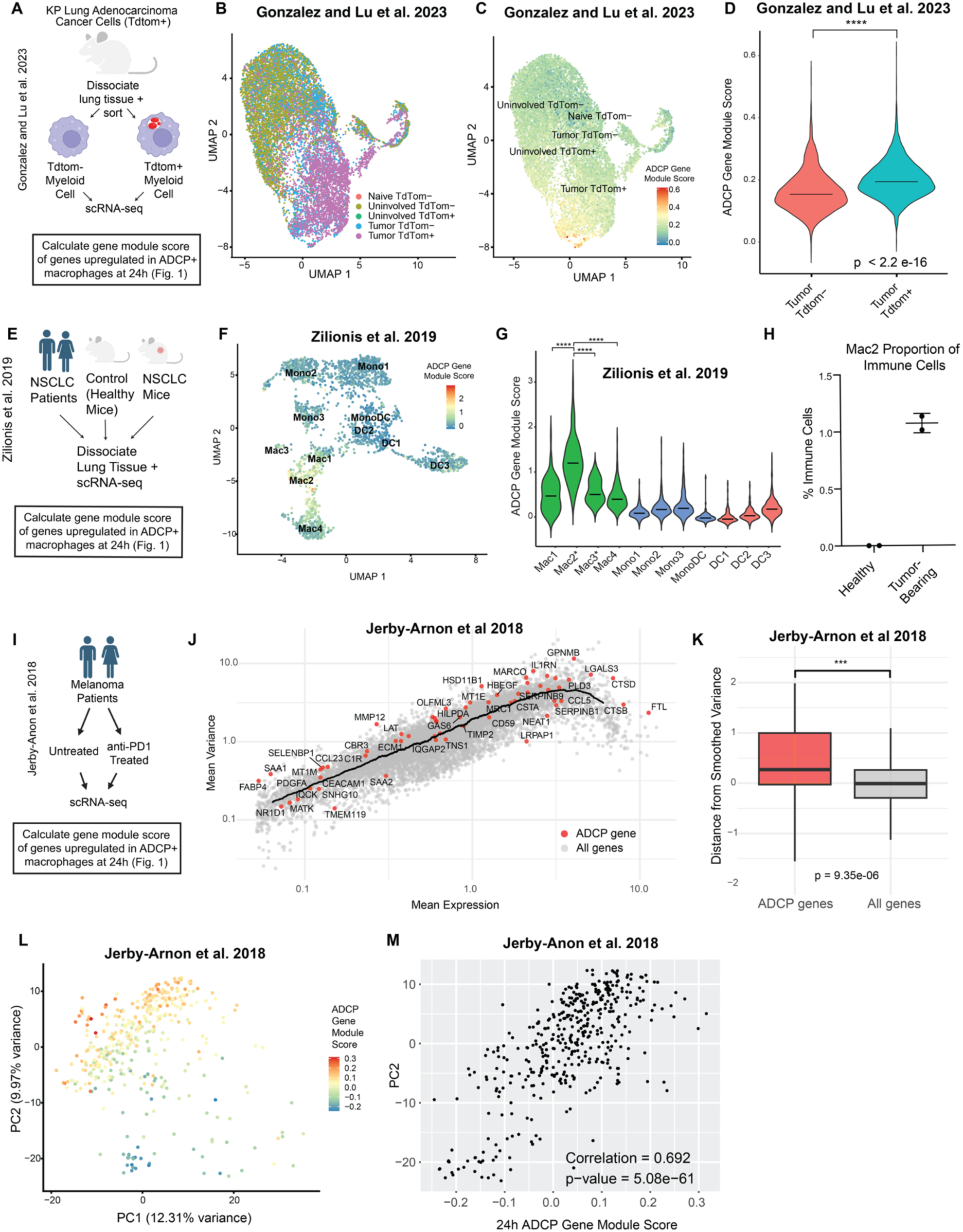
ADCP gene signature is expressed in a tumor-specific macrophage subset and contributes to variance in TAMs *in vivo*. a. Schematic of calculating ADCP gene module score for TdTom+ and TdTom-*in vivo* sorted by phagocytosis of TdTom+ neoplastic cells (Gonzalez and Lu et al. 2023). b. UMAP of *in vivo* alveolar macrophages sorted by tissue TdTom status (Gonzalez and Lu et al. 2023). c. UMAP depicting ADCP gene module score for *in vivo* alveolar macrophages subsets sorted by phagocytosis of TdTom+ neoplastic cells (Gonzalez and Lu et al. 2023). d. Violin plots depicting ADCP gene module score for TdTom+ and TdTom-in vivo alveolar macrophages sorted by phagocytosis of TdTom+ neoplastic cells (Gonzalez and Lu et al. 2023) Statistical significance was determined using the Wilcoxon rank-sum test. e. Schematic of calculating ADCP gene module score for in vivo myeloid cells from lung tissue in NSCLC tumor-bearing and healthy control mice (Zilionis et al. 2019). f. UMAP depicting ADCP gene module score for *in vivo* myeloid subsets from lung tissue in NSCLC tumor-bearing and healthy control mice (Zilionis et al. 2019). g. Violin plots depicting ADCP gene module score for *in vivo* myeloid subsets from lung tissue in NSCLC tumor-bearing and healthy control mice (Zilionis et al. 2019). Statistical significance was determined using the Wilcoxon rank-sum test. h. Proportion of Mac2 subset in immune cells from tumor-bearing and healthy mice (Zilionis et al. 2019). i. Schematic of calculating ADCP gene module score in macrophages from melanoma patients (Jerby-Anon et al. 2018). j. Scatter plot depicting mean variance and mean expression of genes in macrophages from melanoma patients (Jerby-Anon et al. 2018). Line represents running mean variance. k. Box plot depicting distance from smoothed variance of genes in macrophages from melanoma patients (Jerby-Anon et al. 2018). Statistical significance was determined using the Wilcoxon rank-sum test. l. Principal component analysis (PCA) of gene expression profiles of macrophages from melanoma patients (Jerby-Anon et al. 2018). Points are colored by ADCP gene module score. m. Scatter plot depicting PC2 score and ADCP gene module score of macrophages from melanoma patients (Jerby-Anon et al. 2018).

While numerous datasets have mapped the transcriptional heterogeneity of human tumor-associated macrophages *in vivo,*^26–28,101^ the functional characterization of distinct macrophage subsets *in vivo* remains incomplete. To determine if the macrophage heterogeneity observed in other *in vivo* datasets might be partly explained by gene expression programs we observe to be induced by cancer cell phagocytosis, we first used an existing single-cell RNA-seq dataset that profiled myeloid populations in both mouse models and human patients of NSCLC^27^ and calculated the ADCP gene module score in every myeloid cell (Fig. 5e). We observed an enrichment of the ADCP gene module in Mac2 cells (Fig. 5f-g), a macrophage subset characterized by high expression of *Gpnmb, Spp1*, and *Fabp5*.^27^ These Mac2 marker genes were also upregulated in macrophages in our *in vitro* system following ADCP (Fig. 1f). Notably, Mac2 cells were only present in tumor-bearing mice but not in healthy mice (Fig. 5h), suggesting that our ADCP gene signature is expressed by a tumor-specific macrophage subset. We were interested in which gene expression programs were enriched in Mac2 compared to the only other tumor-specific macrophage subset Mac3, and we observed that genes upregulated in Mac2 cells were enriched for lysosomal compartment genes (Fig. S4b-c), which we also observe are upregulated in ADCP+ macrophages (Fig. 1d), suggesting that Mac2 cells may have undergone cancer cell phagocytosis.

We next asked if gene expression programs induced by cancer cell phagocytosis contribute to the observed gene expression variance in human TAMs across multiple cancer types. In a scRNA-seq dataset profiling macrophages in melanoma patients before and after anti-PD1 treatment (Fig. 5i), the variance in expression of ADCP genes across cells in the macrophage compartment was higher on average compared to all genes (Fig. 5j-k). We calculated the ADCP gene module score in macrophages, and in PCA, we observed a significant correlation between ADCP gene module score and PC2, which contributes to 9.97% of the variance (Fig. 5l-m, Fig. S4d). The ADCP gene module score was not differentially expressed upon anti-PD1 treatment (Fig. S4e). We repeated this analysis with macrophages from glioblastoma and synovial sarcoma patients and observed a significant correlation between ADCP gene module score and a top principal component (PC3 in both cases) (Fig. S4f-m).

Collectively, our comparisons to mouse and human *in vivo* scRNA-seq datasets demonstrate that our ADCP gene signature is enriched in phagocytic TAMs *in vivo,* enriched in a tumor-specific macrophage subset *in vivo,* and contributes to observed gene expression variance in human TAMs across multiple cancer types. These findings suggest that cancer cell phagocytosis is a source of TAM heterogeneity and could underlie pro-tumor macrophage transcriptional states *in vivo* even in the absence of mAb treatment.

## Discussion

Here, we profile gene expression and chromatin accessibility changes in macrophages over time following ADCP of live mAb-opsonized cancer cells, compared to changes induced by efferocytosis of apoptotic cancer cells. By comparing the effect of ADCP and efferocytosis in the same *in vitro* system, we identify a shared anti-inflammatory gene expression program and cellular, lysosomal, and oxidative stress TF response in macrophages induced by ADCP and efferocytosis, as well as substrate-specific gene expression changes. ADCP, but not efferocytosis, alters the secretome of macrophages, and conditioned media from ADCP macrophages leads to upregulation of immunosuppressive chemokines in cancer cells and lower expression of EMT genes compared to other macrophage conditioned media. We provide evidence that the ADCP gene signature we observed *in vitro* is expressed in tumor-associated macrophages *in vivo* across numerous cancer types.

We find that following ADCP, TFEB not only activates lysosomal genes but may also unexpectedly activate immune effector genes. While TFEB’s role in activating lysosomal biogenesis genes upon stress has been well established,^102–104^ and its activation of lysosomal proteins upon FcγR-mediated phagocytosis has been demonstrated,^40^ recent studies support a possible broader function of TFEB in innate immunity. TFEB is activated in murine macrophages during *S. aureus* infection and promotes transcription of proinflammatory cytokines, including IL-1β.^105^ Moreover, TFEB- and TFE3-deficient macrophages show impaired transcription and secretion of factors related to differentiation, migration, and inflammation.^78^ Our findings extend this role of TFEB, demonstrating that TFEB may induce leukocyte chemotaxis genes and negative regulators of immune function after ADCP, highlighting its potential dual role in regulating the lysosomal and immune response in macrophages following phagocytosis. Notably, we also observe that TFEB may regulate genes that respond to interleukin-1, including *Il1rn*, an IL-1 receptor agonist that competitively binds IL1R to prevent IL-1α and IL-1β from exerting pro-inflammatory effects. How TFEB activates both pro-inflammatory cytokine expression following bacterial infection and negative regulators of inflammation following ADCP or efferocytosis remains to be explored.

We found that *Gpnmb* expression is strongly induced by both ADCP and efferocytosis in macrophages and that *Gpnmb* transcriptional activation following ADCP is preceded by increased chromatin accessibility at the gene locus. GPNMB+ myeloid cells have been observed across numerous cancer types, and are characterized by expression of immunosuppressive and lipid metabolism genes.^22,64,84,106,107^ In glioblastoma, GPNMB+ SPP1+ macrophages are spatially negatively correlated with CD8+ T cells and associated with upregulated expression of immunosuppressive cytokines and chemokines.^106^ Although GPNMB is highly expressed in TREM2+ and SPP1+ macrophage subsets, which have been associated with poor patient outcomes,^49^ its regulation remains unknown. Although TFEB has been proposed to regulate *Gpnmb* transcriptionally, using our deep learning strategy to identify sequences within regulatory elements which are predictive of accessibility, we did not observe a predictive TFEB family motif within 250kb of the *Gpnm*b locus. Instead, we nominated NFE2L1 and NFE2L2 (NRF2) as potential activators of *Gpnmb* following ADCP, driving increased chromatin accessibility at the locus and increased *Gpnmb* expression. Consistent with this hypothesis, NRF2 deletion in mice led to a decrease in immunosuppressive *Gpnmb*-expressing TREM2^hi^ monocyte-derived macrophages and lower PD-L1 expression in TREM2^hi^ monocyte-derived macrophages.^64^

We also observed an enrichment for NFE family motifs in more accessible elements in macrophages following both ADCP and efferocytosis. Previous studies have demonstrated that NFE2L2 (NRF2) activates antioxidant genes such as *Hmox1* and *Ftl* to promote cellular adaptation to oxidative stress.^60,62^ Consistent with this, we observe upregulation of *Hmox1* in ADCP+ macrophages, suggesting that NRF2 may regulate oxidative stress following phagocytosis. In the context of the TME, NRF2 deletion reduces the immunosuppressive phenotype of tumor-associated macrophages,^64^ suggesting that NRF2 may play a dual role in regulating oxidative stress as well as activating expression of immunosuppressive cytokines.

Future work perturbing the transcriptional activators of ADCP gene expression programs nominated in this study will help identify causal drivers of the anti-inflammatory gene expression program after ADCP and provide insight into how to mitigate the resulting pro-tumor phenotypes and overcome macrophage-intrinsic therapeutic resistance to mAb treatment.

We demonstrate that ADCP, but not efferocytosis, attenuates macrophage-induced expression of EMT genes in cancer spheroids and induces expression of immunosuppressive chemokines in cancer cell spheroids such as *Ccl2, Ccl6*, and *Ccl7* that have been previously shown to recruit macrophages, MDSCs, and regulatory T cells.^98,99^ Future work to identify which secreted factors from ADCP macrophages are responsible for attenuating the pro-EMT phenotype can expand our understanding of how macrophages promote EMT in cancer cells and how cancer cell phagocytosis alters the macrophage secretome. Additionally, consistent with recent reports,^94^ we demonstrate that efferocytosis, but not ADCP, uniquely induces upregulation of *Egr1* and increased activity of EGR family TFs.

While our *in vitro* experimental system allowed us to directly probe time- and substrate-dependent effects of cancer cell phagocytosis on macrophages, macrophages in the TME are heterogeneous and subject to a more complex milieu of cell signals. For example, previous studies have demonstrated that efferocytosis combined with exposure to secreted factors led to distinct transcriptional changes in macrophages not observed after efferocytosis alone,^9^ and macrophages *in vivo* are exposed to numerous secreted factors that could modulate the effect of phagocytosis. Furthermore, we used LPS as a model to polarize macrophages towards a pro-inflammatory phagocytic state, although many pro-inflammatory stimuli including IFNγ exist in the TME. Underlying heterogeneity in macrophages from lineage, tissue of origin, and initial polarization state may interact with distinct types of phagocytosis to induce molecular changes. While many of these parameters remain to be explored in future work, we demonstrate that the ADCP gene signature we observed in our highly controlled model system *in vitro* is expressed in tumor-associated macrophages *in vivo* across numerous cancer types in both model systems and patient settings. We observed higher expression of our ADCP gene module in phagocytic TAMs *in vivo,* and observed an enrichment for our ADCP gene signature in a tumor-specific macrophage subset *in vivo*.

Numerous studies have profiled the transcriptional and epigenetic heterogeneity of tumor-associated macrophages *in vivo;* however, the mechanisms by which pro-tumor macrophage states arise and the contribution of prior phagocytic activity to these molecular states has been unclear. We demonstrate that ADCP induces increased activity by cellular, lysosomal, and oxidative TFs as well as a pro-angiogenic and immunosuppressive cytokine gene expression program, observed *in vitro* and in TAMs *in vivo.* Overall, these findings expand our understanding of how cancer cell phagocytosis contributes to pro-tumor macrophage molecular states, and set the stage for future work to unravel how pro-tumor states contribute to resistance and non-response to mAb treatment.

## Acknowledgements

We thank Peter Du and Katherine Liu for their advice on scRNA-seq analysis, Betty Liu for advice on ATAC-seq analysis, Soon Il Higashino for keeping our laboratory running, and members of the Bassik and Greenleaf labs for helpful discussions. This work was supported by a NSF Graduate Fellowship (C.R.Z.), and NCI U54CA261719 (M.C.B). W.J.G acknowledges support from the Arc Institute and the Chan-Zuckerberg Biohub.

## Competing Interests

W.J.G. is a consultant and equity holder for 10x Genomics, Guardant Health, Quantapore, and Ultima Genomics and cofounder of Protillion Biosciences and is named on patents describing ATAC-seq. A.K. is on the scientific advisory board of SerImmune, TensorBio, AINovo, is a consultant with Arcadia Science, Inari, Precede Biosciences, was a consultant with Illumina and PatchBio and has a financial stake in DeepGenomics, Immunai and Freenome. M.C.B declares outside interest in DEM Biopharma and Stylus Medicine. All other authors declare no competing interests.

## Author Contributions

C.R.Z., M.C.B, and W.J.G conceived of the study. C.R.Z. performed experiments, next-generation sequencing, and analysis of RNA-seq, ATAC-seq, and scRNA-seq data. M.G. and C.R.Z. collected conditioned media and conducted phagocytosis assays for the cancer cell spheroid experiments. L.X. trained ChromBPNet models and performed analysis related to correlation between contribution scores and RNA expression with input from A.K., W.J.G., and C.R.Z. C.R.Z., M.C.B., and W.J.G wrote the manuscript, with input from all authors. M.C.B. and W.J.G. jointly supervised the work.

## Data Availability

RNA-seq and ATAC-seq data that support the findings of this study have been deposited in SRA with the BioProject PRJNA1262986.

## Code Availability

Our use of published software tools, with description of their use and parameter settings, is available in Methods.

## Supplementary Text

Clusters 3-5 were primarily differentially expressed over time, but not differentially expressed between ADCP+ and ADCP-macrophages. Cluster 3 (*Cd80*+ *Cd86*+) genes were more highly expressed at early time points and enriched for genes in cytokine production and response to LPS pathways, consistent with the effect of washing out the initial LPS stimulus. Cluster 4 (*Ki67+ Top2a*+) genes were expressed more highly at later time points and enriched for cell division genes. Cluster 5 (*Il10*+ *Arg1*+), which was composed of genes that increased in expression over time with co-culture with cancer cells and after the initial LPS stimulation was washed out, was also enriched for genes in angiogenesis and response to wounding pathways. Overall, these gene clusters were characterized by increasing expression of pro-inflammatory LPS-response genes and decreasing expression of anti-inflammatory genes over time.

**Supplementary Figure 1.**
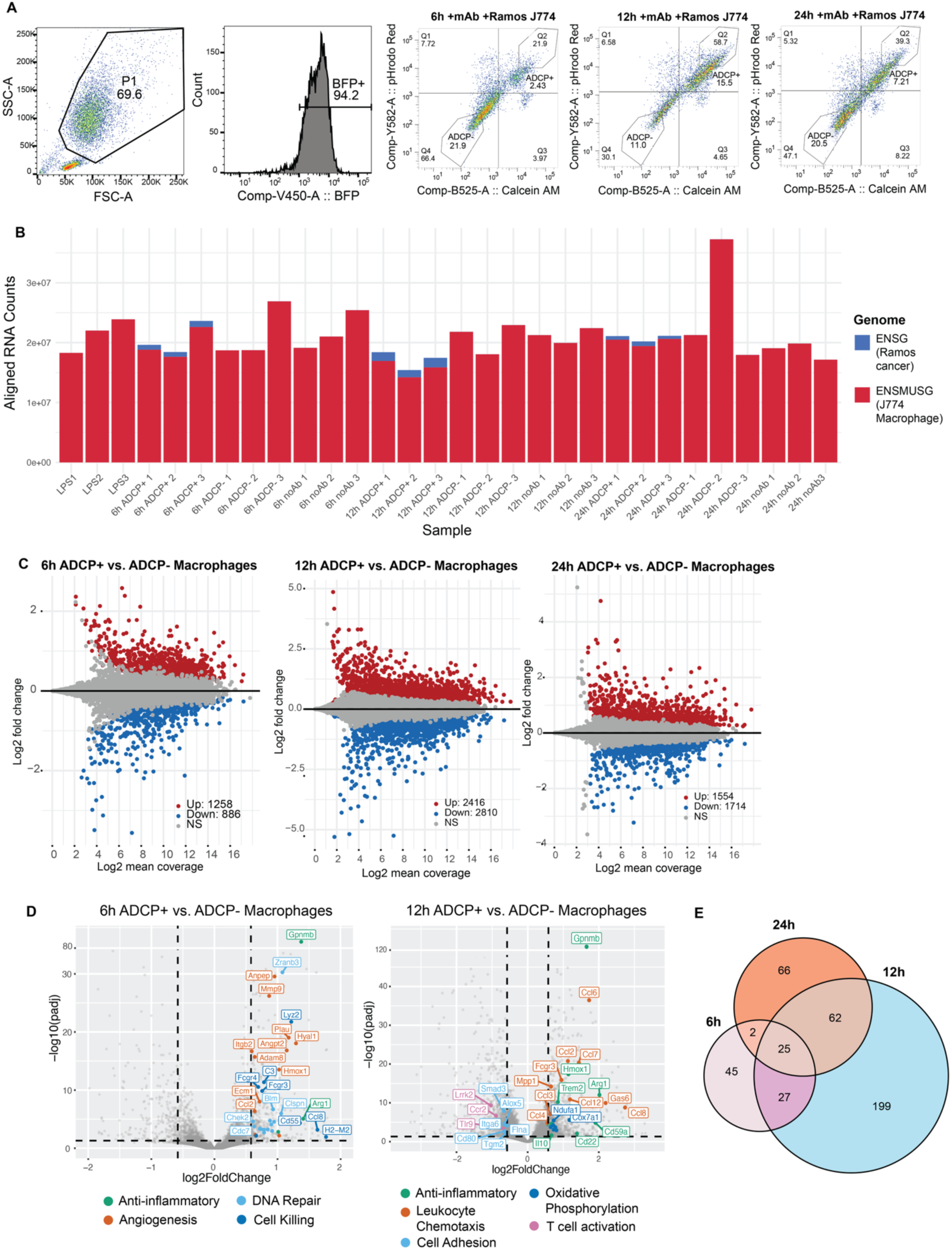
Profiling gene expression changes after ADCP over time in macrophages. a. FACS gating on ADCP live cancer cell phagocytosis time course. Cells were first gated on BFP+ signal to sort J774 macrophages. In the presence of mAb, pHrodo+ calcein+ cells were sorted as ADCP+ macrophages and pHrodo-calcein-macrophages were sorted as ADCP-macrophages. b. Aligned RNA reads across samples. Bars are colored based on genome alignment (ENMUSG counts originate from murine J774 macrophages while ENSG counts originate from human Ramos lymphoma cells). c. Differential expression between ADCP+ and ADCP-macrophages at each time point. The x-axis represents the log2 mean reads per gene and the y-axis represents the log2 fold change in expression. Significantly higher expressed genes (log2FC > 1, adjusted p-value < 0.05) are colored in red, and significantly lower expressed genes (log2FC < -1, adjusted p-value < 0.05) are colored in blue. d. Differential expression between ADCP+ and ADCP-macrophages plotted at each time point. The x-axis represents the log2 fold change in expression and the y-axis represents the adjusted p-value. Genes are colored by enriched GO terms. e. Venn diagram of upregulated genes between ADCP+ and ADCP-macrophages at each time point.

**Supplementary Figure 2.**
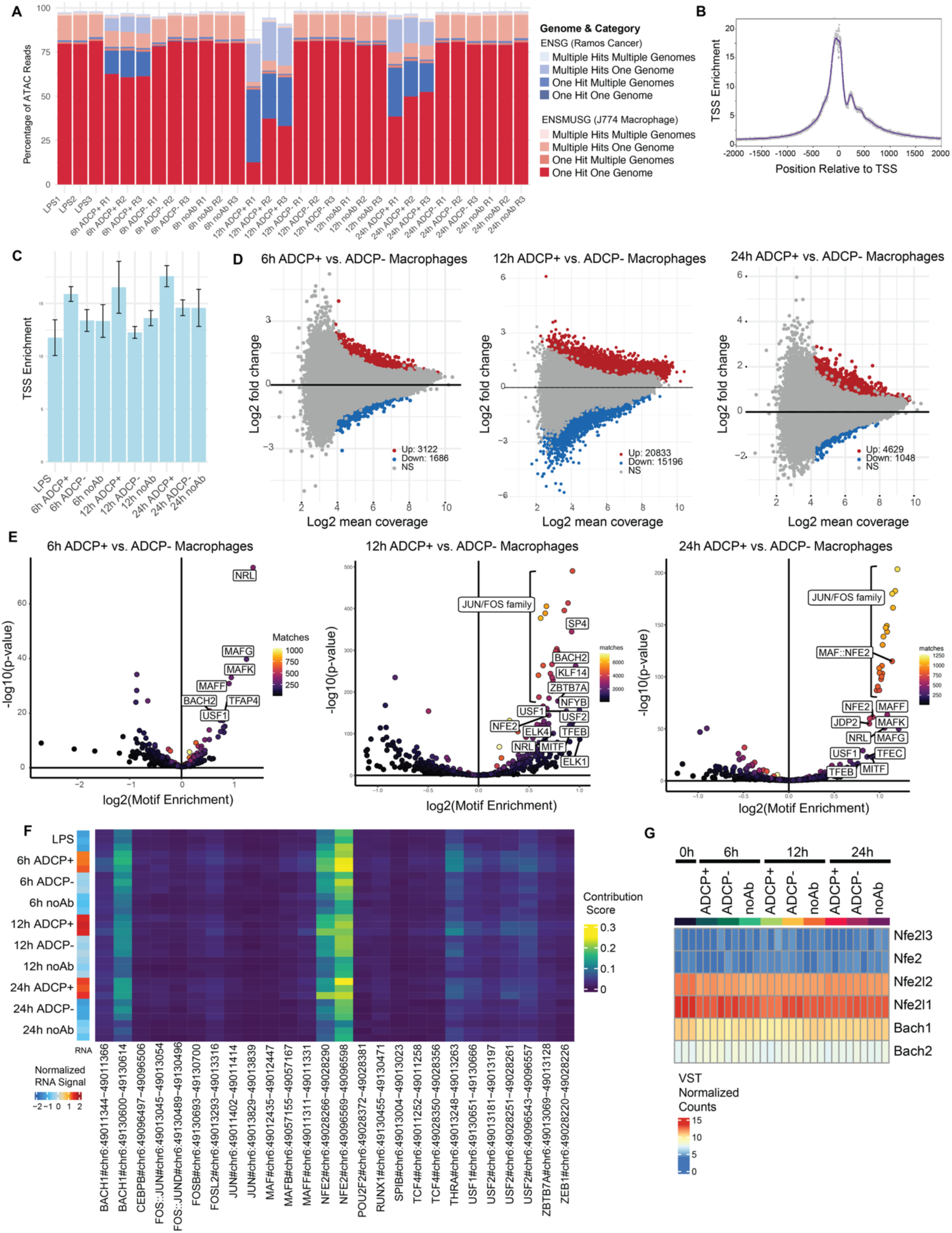
Profiling chromatin accessibility changes after ADCP over time in macrophages. a. Aligned ATAC reads across samples. Bars are colored based on genome alignment and number of hits (ENMUSG counts originate from murine J774 macrophages while ENSG counts originate from human Ramos lymphoma cells). b. Representative plot of aggregate signal around the transcription start site (TSS) for all ATAC-seq peaks in one ADCP+ macrophage sample. This plot represents the signal-to-noise quantification of our ATAC-seq data. TSS enrichment scores greater than 10 indicate high quality ATAC-seq data. c. TSS enrichment scores for ATAC-seq libraries. d. Differential accessibility across ATAC-seq peaks between ADCP+ and ADCP-macrophages at each time point. The x-axis represents the log2 mean accessibility per peak and the y-axis represents the log2 fold change in accessibility. Significantly more accessible peaks (log2FC > 0.58, adjusted p-value < 0.05) are colored in red, and significantly less accessible peaks (log2FC < -0.58, adjusted p-value < 0.05) are colored in blue. e. TF motif enrichment in more accessible peaks in ADCP+ vs. ADCP-macrophages at each time point. f. ChromBPNet model-derived contribution scores at TF motifs that are highly correlated (Pearson correlation > 0.5) with Gpnmb RNA expression within 100 kb of GPNMB TSS. g. RNA expression of BACH/NFE transcription factor family members across samples. Heatmap represents VST normalized counts.

**Supplementary Figure 3.**
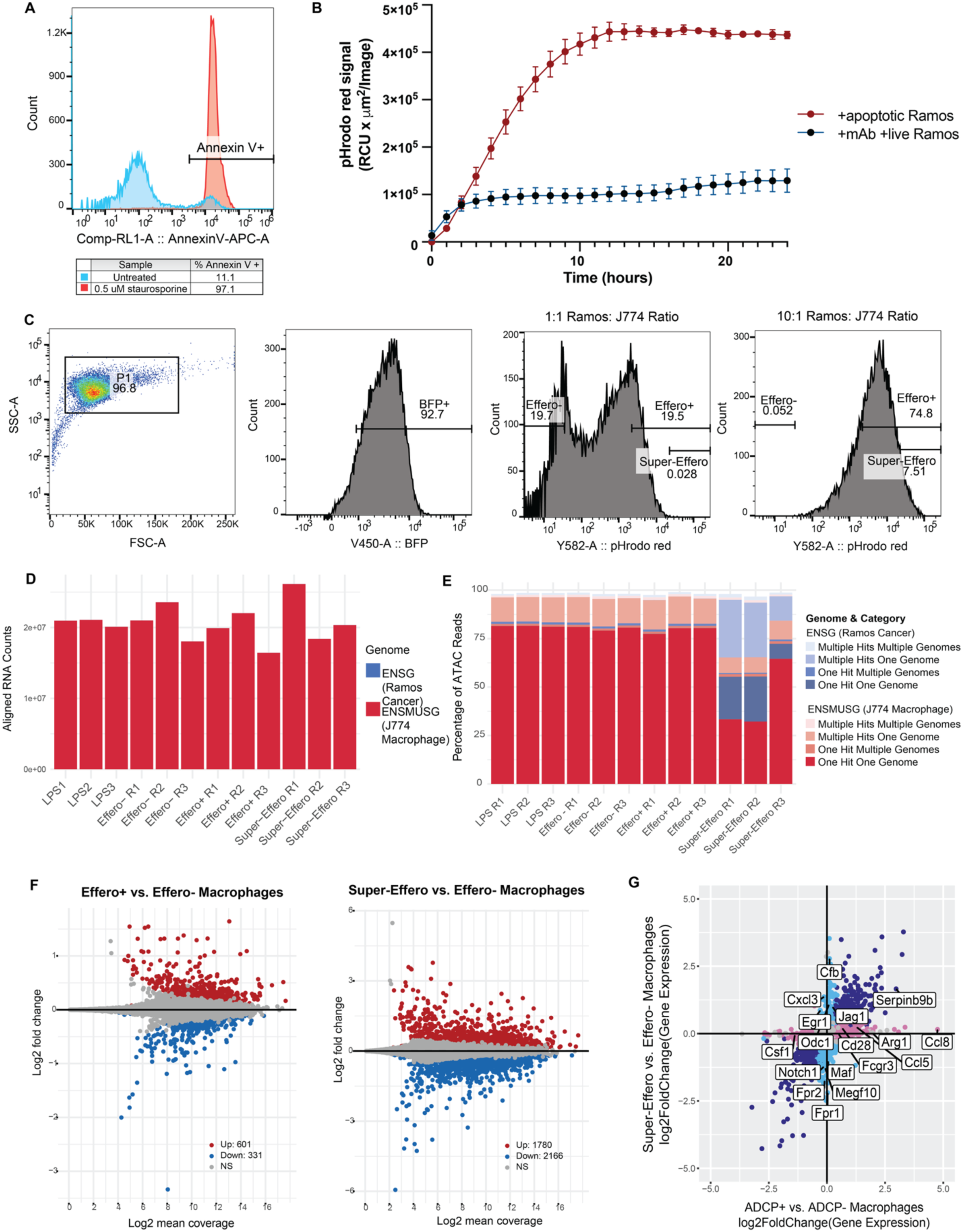
Profiling gene regulatory changes following efferocytosis in macrophages. a. Annexin V staining of staurosporine-treated and untreated Ramos cells. b. Phagocytosis assay quantifying pHrodo red signal (which increases upon acidification in the lysosome) in macrophages over time cultured with either mAb-opsonized live Ramos cells or apoptotic Ramos cells. Points are colored based on phagocytic substrate. c. FACS gating on ADCP live cancer cell phagocytosis time course. Cells were first gated on BFP+ signal to sort J774 macrophages. At a low co-culture ratio of 1:1 apoptotic Ramos:J774, pHrodo red+ cells were sorted as Effero+ macrophages and pHrodo-cells were sorted as Effero-macrophages. At a high co-culture ratio of 10:1 apoptotic Ramos:J774, the top 10% of pHrodo red+ cells were sorted as Super-Effero macrophages. d. Aligned RNA reads across samples. Bars are colored based on genome alignment (ENMUSG counts originate from murine J774 macrophages while ENSG counts originate from human Ramos lymphoma cells). e. Aligned ATAC reads across samples. Bars are colored based on genome alignment and number of hits (ENMUSG counts originate from murine J774 macrophages while ENSG counts originate from human Ramos lymphoma cells). f. Differential expression between Effero+ and Effero-macrophages, and Super-Effero and Effero-macrophages. The x-axis represents the log2 mean reads per gene and the y-axis represents the log2 fold change in expression. Significantly higher expressed genes (log2FC > 1, adjusted p-value < 0.05) are colored in red, and significantly lower expressed genes (log2FC < -1, adjusted p-value < 0.05) are colored in blue. g. TF motif enrichment in more accessible peaks (n=3 biological replicates per substrate type) upon ADCP (ADCP+ vs. ADCP-log2Enrichment) or higher amounts of efferocytosis (Super-Effero vs. Effero-log2Enrichment). Colored points represent significant TF motif enrichment (p-adj < 0.05).

**Supplementary Figure 4.**
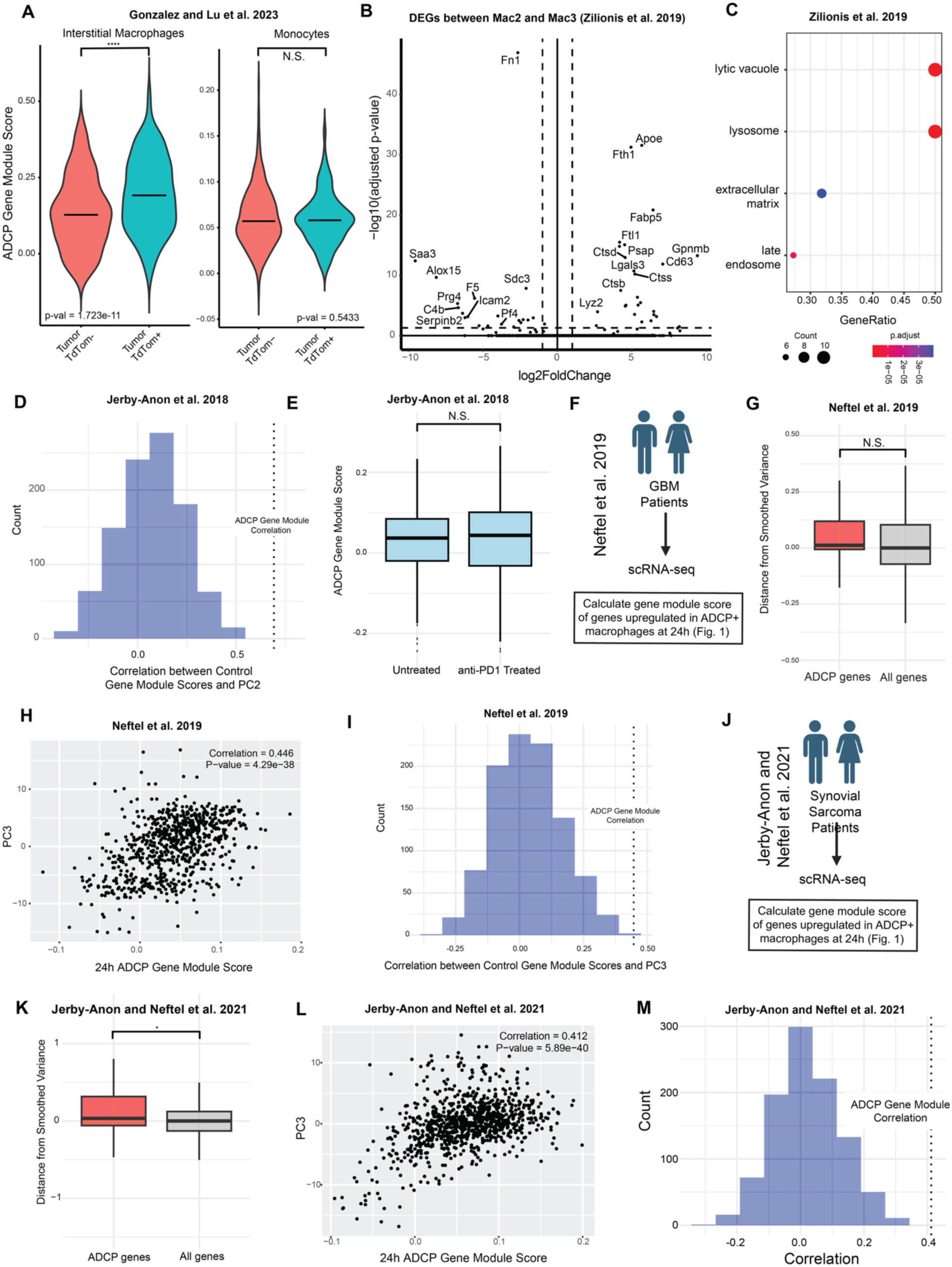
ADCP gene signature is expressed in TAMs *in vivo*. a. ADCP gene module score for TdTom+ and TdTom-*in vivo* interstitial macrophages and monocytes sorted by phagocytosis of TdTom+ neoplastic cells.^23^ Statistical significance was determined using the Wilcoxon rank-sum test. b. Differentially expressed genes between Mac2 and Mac3 subsets.^27^ c. GO term enrichment in upregulated genes in Mac2 compared to Mac3 cells.^27^ d. ADCP gene module score in macrophages from untreated and anti-PD1 treated patients.^108^ e. Distribution of correlations between PC2^108^ and gene module scores of 1000 control gene sets of randomly selected genes with similar expression to ADCP genes. Dotted line represents the correlation between ADCP gene module score and PC2. f. Schematic of calculating ADCP gene module score for macrophages from glioblastoma patients.^109^ g. Box plot depicting distance from smoothed variance of genes in macrophages from glioblastoma patients.^109^ h. Scatter plot depicting PC2 score^109^ and ADCP gene module score of macrophages from glioblastoma patients with Pearson correlation value. i. Distribution of correlations between PC2^109^ and gene module scores of 1000 control gene sets of randomly selected genes with similar expression and variance to ADCP genes. Dotted line represents the correlation between ADCP gene module score and PC2. j. Schematic of calculating ADCP gene module score for macrophages from synovial sarcoma patients.^110^ k. Box plot depicting distance from smoothed variance of genes in macrophages from synovial sarcoma patients.^110^ l. Scatter plot depicting PC2 score^110^ and ADCP gene module score of macrophages from glioblastoma patients with Pearson correlation value. m. Distribution of correlations between PC2^110^ and gene module scores of 1000 control gene sets of randomly selected genes with similar expression and variance to ADCP genes. Dotted line represents the correlation between ADCP gene module score and PC2.

## Supplementary Tables

**Supplementary Table 1**. DEGs between ADCP+ and ADCP-macrophages at each time point (6h, 12h, 24h)

**Supplementary Table 2.** K-means clustering on macrophage gene expression profiles following ADCP

**Supplementary Table 3**. Motif Enrichment in differentially accessible peaks between ADCP+ and ADCP-macrophages at each time point (6h, 12h, 24h)

**Supplementary Table 4.** Fischer’s exact test of TF family motif enrichment in nearby peaks to differentially expressed gene programs (12h, 24h)

**Supplementary Table 5.** ChromBPNet model-derived contribution scores and correlation to gene expression at motifs within 100kb of *Gpnmb* TSS

**Supplementary Table 6.** DEGs between ADCP+ vs Effero+, ADCP+ vs Super-Effero, and Effero+ vs. Super-Effero macrophages

**Supplementary Table 7.** Motif enrichment in differentially accessible peaks between ADCP+ vs Effero+, ADCP+ vs Super-Effero, and Effero+ vs. Super-Effero macrophages

**Supplementary Table 8.** DEGs between cancer cell spheroids seeded in ADCP+ macrophage conditioned media and ADCP-macrophage conditioned media

## Methods

### Sorting based on antibody-dependent cellular phagocytosis for time course

J774 cells were lifted and plated in 15 cm plates at a density of 5 million cells per plate in 30 mL of medium 48h before the start of the experiment. At 24 h after plating, medium was aspirated and replaced with fresh medium containing 100 ng ml^−1^ LPS (Sigma). On the day of the experiment, target Ramos cells were counted and incubated in PBS containing 100 nM pHrodo-Red succinimidyl ester (Thermo Fisher) and 50 nM calcein AM (Invitrogen) at a cell concentration of 1 million ml^−1^ for 15 min at 37 °C, pelleted, and resuspended in DMEM supplemented with 2 mM glutamine, 100 U ml^−1^ penicillin, 100 µg ml^−1^ streptomycin and 10% heat-inactivated FBS, containing 10 𝞵g/mL anti-CD47 (clone B6.H12, BioXCell) and 15 𝞵g/mL anti-CD20 antibodies (Rituxan) at a cell concentration of 1 × 10^6^ ml^−1^ at a 10:1 ratio of Ramos:J774. To control for the effect of co-culture with Ramos cells, pHrodo-and calcein-labeled target Ramos cells were also added to J774 cells in the absence of anti-CD20 and anti-CD47 at a 10:1 ratio of Ramos:J774. Target Ramos cells were co-cultured with J774 cells for either 6, 12, or 24 hours. For the time course, J774 cells were stimulated with LPS and co-cultured with target Ramos cells in a reverse time course such that sorting for all time points occurred synchronously.

After co-culture with Ramos cells, J774 cells were washed 3 times with warm PBS to detach any bound un-internalized Ramos cells. J774 cells were lifted by incubating with 10 mL accutase for 10 min and gentle scraping. J774 cells were washed once in medium and resuspended in fresh medium and filtered before sorting. Cells were gated on BFP+ cells (J774s were transduced with an empty sgRNA BFP plasmid) to differentiate from un-internalized Ramos cells. J774 cells were sorted into live cell eaters (pHrodo red+ calcein+) and live cell non-eaters (pHrodo red-calcein-). 300,000 cells were sorted per condition. After FACS, sorted cells were washed in medium and ¼ (75,000) of the cells were aliquoted for ATAC-seq and ¾ of the cells (225,000) were snap frozen for RNA-seq. Phagocytosis assays were done in triplicate, and samples were sorted separately before downstream RNA-seq or ATAC-seq.

### Sorting based on apoptotic cancer cell phagocytosis

J774 cells were lifted and plated in 15 cm plates at a density of 5 million cells per plate in 30 mL of medium 48h before the start of the experiment. At 24 h after plating, medium was aspirated and replaced with fresh medium containing 100 ng ml^−1^ LPS (Sigma). Ramos cells were treated with 0.5 uM staurosporine (Sigma-Aldrich) to induce apoptosis 24h before the start of the experiment. On the day of the experiment, apoptotic target Ramos cells were counted and incubated in PBS containing 100 nM pHrodo-Red succinimidyl ester (Thermo Fisher) for 15 minutes. pHrodo labeled cells were washed in DMEM and added at either a high ratio of 10:1 Ramos:J774 (to sort Super-Effero macrophages) or low ratio of 1:1 Ramos:J774 (to sort Effero+ and Effero-macrophages). After 24h after co-culture with apoptotic Ramos cells, J774s were washed 3 times with warm PBS to detach any bound unengulfed Ramos cells. J774 cells were lifted by incubating with 10 mL accutase for 10 min and gentle scraping. J774 cells were washed once in medium and resuspended in fresh medium and filtered before sorting. Cells were gated on BFP+ cells (J774s were transduced with an empty sgRNA BFP plasmid) to differentiate from uninternalized Ramos cells. At the low co-culture ratio, BFP+ J774 cells were gated on pHrodo red signal into pHrodo red+ cells (Effero+ macrophages) and pHrodo red-cells (Effero-macrophages). At the high co-culture ratio, BFP+ J774 cells were gated on the top 10% of pHrodo red signal and sorted as apoptotic super-eaterse. 300,000 cells were sorted per condition. After FACS, sorted cells were washed in medium and ¼ (75,000) of the cells were aliquoted for ATAC-seq and ¾ of the cells (225,000) were snap frozen for RNA-seq. Phagocytosis assays were done in triplicate, and samples were sorted separately before downstream RNA-seq or ATAC-seq.

### Time lapse microscopy assay for phagocytosis

J774 cells were lifted by scraping, counted, and plated in 24-well tissue culture plates at a density of 75,000 cells per well in 0.5 ml medium 48 h before the start of the experiment. At 24 h after plating, medium was aspirated and replaced with fresh medium containing 100 ng ml−1 LPS (Sigma). On the day of the experiment, target Ramos cells were counted and incubated in PBS containing 100 nM pHrodo-Red succinimidyl ester (Thermo Fisher) at a cell concentration of 1 million ml−1 for 15 min at 37 °C, pelleted, and resuspended in DMEM supplemented with 2 mM glutamine, 100 U ml−1 penicillin, 100 µg ml−1 streptomycin and 10% heat-inactivated FBS, and containing anti-CD20 and anti-CD47 antibodies, as indicated, at a cell concentration of 1 × 10^6^ ml^−1^. Plates were transferred to an incubator and imaged every 60 min using an Incucyte (Essen). Total red intensity for each well, averaged over 9–16 images per well, was calculated after applying a threshold for red intensity that excluded most or all cells at the first time point using top-hat background subtraction. Reported values represent the mean total red fluorescence intensity at each timepoint. Phagocytosis assays were done in 4 biological replicates.

### Cell culture

Ramos cells were maintained in suspension culture in RPMI-1640 supplemented with 2 mM glutamine, 100 U ml−1 penicillin, 100 µg ml−1 streptomycin and 10% heat-inactivated fetal bovine serum (FBS). J774 cells were cultured in DMEM supplemented with 2 mM glutamine, 100 U ml−1 penicillin, 100 µg ml−1 streptomycin and 10% heat-inactivated FBS and were passaged when nearing confluency using Accutase and scraping. 802T4 cells, a mouse lung cancer cell line derived from a primary lung tumor in a Kras^LSL-G12D/+^;p53^flox/flox^ mouse model, were obtained from the Winslow Lab at Stanford ^111^. 802T4 cells were cultured in DMEM supplemented with 2 mM glutamine, 100 U ml−1 penicillin, 100 µg ml−1 streptomycin and 10% heat-inactivated FBS and were passaged when nearing confluency using Trypsin.

### Bulk ATAC-seq

Approximately 75,000 cells were resuspended in 250 μl PBS and centrifuged at 500*g* for 5 min at 4 °C in a fixed-angle centrifuge. Pelleted cells were resuspended in 50 μl ATAC–seq resuspension buffer (RSB; 10 mM Tris-HCl pH 7.4, 10 mM NaCl and 3 mM MgCl_2_ in ddH_2_O made fresh) containing 0.1% NP40, 0.1% Tween-20 and 0.01% digitonin according to the omni-ATAC–seq protocol. After incubating on ice for 3 min, 1 ml of ATAC–seq RSB containing 0.1% Tween-20 was added. Nuclei were centrifuged at 500*g* for 5 min at 4 °C in a fixed-angle centrifuge, 900 μl of the supernatant was taken off, and the nuclei were centrifuged for an additional 5 min under the same conditions. The remaining 200 μl of supernatant was aspirated and nuclei were resuspended in 50 μl of transposition mix (25 μl 2× TD buffer (2 ml 1 M Tris-HCl, pH 7.6, 1 ml 1 M MgCl2, 20 ml DMF and 77 ml ddH_2_O aliquoted and stored at −20 °C), 2.5 μl transposase (100 nM final), 16.5 μl PBS, 0.5 μl 1% digitonin, 0.5 μl 10% Tween-20 and μl ddH_2_O). Transposition reactions were incubated at 37 °C for 30 min with 1,000 rpm shaking in a thermomixer and cleaned up using MinElute PCR purification columns (Qiagen). The transposed samples were then amplified to add sample indices and sequencing flow-cell adapters and cleaned up with MinElute PCR purification columns (Qiagen), with a target concentration of 20 μl at 4 nM. Paired-end sequencing (2x 36bp) was performed on an Illumina NextSeq using 75 cycle kits. Raw ATAC-seq data has been deposited in SRA with the BioProject PRJNA1262986.

### Bulk RNA-seq

Approximately 200,000 cells were collected for RNA-seq and centrifuged at 500g for 5 min. Supernatant was aspirated and cells were snap frozen in liquid nitrogen. RNA was extracted using Qiagen RNeasy Mini kit (Qiagen 74104). Library preparation, quality control and RNA sequencing was performed by Novogene using Illumina NovaSeq 6000 using a 150 bp paired-end sequencing protocol. Raw RNA-seq data has been deposited in SRA with the BioProject PRJNA1262986.

### RNA-seq preprocessing and analysis

Transcripts were mapped to the ICRG reference genome^112^ using STAR (v2.7.0)^113^ and gene-level counts were generated using HTSeq (v.0.13.5).^114^ Only ENSMUSG counts were used for downstream analysis and followed by differential gene expression analysis using DESeq2 (1.28.1).^115^ GO term enrichment was performed using clusterProfiler (version 3.0.4).

### ATAC-seq preprocessing and analysis

All preprocessing code is available on https://github.com/GreenleafLab/snakeATAC_singularity. Because of the presence of both human (from phagocytosed human Ramos cancer cells) and mouse (from murine J774 macrophage cells) reads, we adapted a metagenomics pipeline (https://www.metagenomics.wiki/tools/short-read/remove-host-sequences) to remove human reads. We first mapped reads using bowtie2 to the hg38 reference genome, keeping both aligned and unaligned reads (paired-end reads) (“bowtie2 --threads {threads} -x ” + REFERENCE_FILE_HUMAN + “ -1 {input.left} -2 {input.right} -S {output}”). We discarded any reads that mapped to the hg38 reference genome (“samtools view -b -f 12 -F 256 {input} > {output}”). We split paired-end reads into separated fastq files (“samtools sort -n -m 5G -@ 2 {input} -o {output}” “samtools fastq -@ 8 {input} -1 {output.left} -2 {output.right} -0 /dev/null -s /dev/null -n”). We then used the output fastq files (which have removed human reads) with the standard snakeATAC pipeline using the mm10 reference genome, TSS coordinates, and ENCODE mm10 blacklist. We used ChrAccR (https://github.com/GreenleafLab/ChrAccR) to perform the ATAC peak calling, PCA, and motif differential analysis.

### Motif enrichment in nearby Peaks to ADCP gene expression programs

At 12 hours and 24 hours, we selected GO terms enriched in upregulated genes (log2FC > 0.58, adjusted p-value < 0.05) and TF family motifs enriched in more accessible peaks (log2FC > 0.58, adjusted p-value < 0.05) in ADCP+ macrophages compared to ADCP-macrophages. For each GO term, we identified all peaks within 250kb of the TSS of each gene. We then calculated motif enrichment using Fisher’s exact test of each TF family motif in more accessible peaks compared to unchanging peaks.

### Preprocessing of ATAC-seq data for ChromBPNet models

We followed the workflow (https://github.com/kundajelab/scAnnot-to-chrombpnet) and prepared a peak set and corresponding base-resolution bigWig file as inputs for each bulk ATAC-seq fragment to train ChromBPNet. Tn5 insertion sites were randomly assigned to two separate pseudoreplicates. These pseudoreplicates, along with a concatenated total-pseudoreplicate file, were used for peak calling with MACS2 (https://github.com/macs3-project/MACS). Peaks were included if those called from the total-pseudoreplicate that overlapped with peaks from both individual pseudoreplicates. The final union peak set, created by merging all retained peaks, was used as input for ChromBPNet model training and downstream analyses.

### Training ChromBPNet models

We trained bias-correction ChromBPNet models (ChromBPNet v0.1.7, https://github.com/kundajelab/chrombpnet) and followed the workflow described by Pampari et al.^80^ using the peaks file and bigWig file generated in the preprocessing of ATAC-seq data. For each sample, we used a five-fold cross-validation scheme to ensure each chromosome appeared in the test set of at least one cross-validation fold.

### Inferring predictive motif instances

To scan motif instances in each sample, we used the FiNeMo v0.15 package (https://github.com/austintwang/finemo_gpu) with lambda regularization parameter alpha of 0.7 and the sequence patterns identified by TF-MoDISco as inputs. All instances of motifs across samples were merged by keeping the instances with highest “hit_correlation” value in cases of an overlap. To quantify differences in TF regulation across samples, we assigned the maximum contribution score across all nucleotides within the motif instance to represent contribution of the motif.

### Linking transcription factors (TFs) to target genes

To identify potential transcription factor (TF) candidates regulating target genes, we computed the Pearson correlation between the contribution scores of predictive motif instances located within ±100 kb of each gene’s transcription start site (TSS) and the corresponding gene expression levels across samples. Contribution scores for each predictive motif instance were averaged across identical annotations. Gene expression levels were normalized to a total of 10,000 counts per experiment, followed by log1p transformation. We retained motif instances with a Pearson correlation above 0.7. Finally, predictive motif instances were mapped to their corresponding TFs to establish TF–gene regulatory linkages.

### Gene expression comparisons between ADCP and efferocytosis

To identify shared and distinct differentially expressed genes upon ADCP and efferocytosis in macrophages, we extracted DE results from our pairwise comparisons between Super-Effero and Effero-macrophages, Effero+ and Effero-macrophages, and ADCP+ and ADCP-macrophages at 24 hours. For each gene, we plotted its fold change between each comparison respectively and colored points by its significance.

### Chromatin accessibility comparisons between ADCP and efferocytosis

To identify shared and distinct TFs driving chromatin accessibility changes upon ADCP and efferocytosis, we extracted the TF motif enrichment results from our pairwise comparisons between Super-Effero and Effero-macrophages, Effero+ and Effero-macrophages, and ADCP+ and ADCP-macrophages at 24 hours. For each TF motif, we plotted its motif enrichment between each comparison respectively and colored points by its significance.

### Cancer cell spheroid culture in macrophage conditioned media and RNA-seq

J774 macrophages were lifted and plated in 6-well plates at a density of 330,000 per well in 3 mL of media 48h before the start of the experiment. At 24 h after plating, medium was aspirated and replaced with fresh medium containing 100 ng ml^−1^ LPS (Sigma). For ADCP macrophages, on the day of the experiment, target Ramos cells were counted and incubated in PBS containing 100 nM pHrodo-Red succinimidyl ester (Thermo Fisher) and 50 nM calcein AM (Invitrogen) at a cell concentration of 1 million ml^−1^ for 15 min at 37 °C, pelleted, and resuspended in DMEM supplemented with 2 mM glutamine, 100 U ml^−1^ penicillin, 100 µg ml^−1^ streptomycin and 10% heat-inactivated FBS, and containing anti-CD20 and anti-CD47 antibodies, as indicated, at a cell concentration of 1 × 10^6^ ml^−1^ at a 10:1 ratio of Ramos:J774 for 48 hours. For macrophages co-cultured with live cancer cells, pHrodo- and calcein-labeled target Ramos cells were also added to J774 cells in the absence of anti-CD20 and anti-CD47 at a 10:1 ratio of Ramos:J774. Target Ramos cells were co-cultured with J774 cells for 48 hours. For efferocytic macrophages, Ramos cells were treated with 0.5 uM staurosporine to induce apoptosis 24h before the start of the experiment. On the day of the experiment, apoptotic target Ramos cells were counted and incubated in PBS containing 100 nM pHrodo red for 15 minutes. pHrodo labeled cells were washed in DMEM and added at a ratio of 10:1 Ramos:J774 for 48 hours. 48 hours after the phagocytosis assay, conditioned media was collected.

To seed 802T4 lung adenocarcinoma spheroids, 802T4 cells were lifted and spun down at 500g for 5 minutes. 802T4 cells were resuspended at a concentration of 200,000 cells/mL and 500 uL of cell solution (100,000 cells) was aliquoted into Eppendorf tubes and spun down at 500g for 5 minutes. 802T4 cell pellets were then resuspended in 500 uL of conditioned media into a single-cell suspension. Untreated spheroids were resuspended in 500 uL of cell culture media. 500 uL of 1.5% methylcellulose (Fisher, no. M-352) was added to each aliquot of conditioned media 802T4 cell solution for a final concentration of 0.75% methylcellulose and plated into a well of a 24-well ultra low-attachment plate. After 7 days of culture in conditioned media, spheroids and methylcellulose media were transferred to Eppendorf tubes. Cells were spun down at 600g for 5 minutes, washed with PBS, and snap frozen. Phagocytosis assays to generate conditioned media were done in triplicate, followed by spheroid seeding and culture.

### Analysis of scRNA-seq datasets

The ADCP gene module was defined as genes with higher expression (log2FC > 1, adjusted p-value < 0.05) in ADCP+ compared to ADCP-macrophages. In the Gonzalez and Lu et al. dataset, the alveolar macrophage compartment, which had the greatest proportion of phagocytic macrophages, was subsetted and used for downstream analysis. The ADCP gene module score was calculated using the Seurat package and a Wilcoxon-rank sum test was used to determine statistical significance of the ADCP gene module score between TdTom+ and TdTom-macrophages. The same analysis was performed on the interstitial macrophage and monocyte compartments.

In the Zilionis et al. dataset, the myeloid compartment was subsetted and used for downstream analysis. The ADCP gene module score was calculated using the Seurat package. For differential expression analysis between Mac2 and Mac3 cells, the two tumor-specific macrophage subsets, raw counts were pseudobulked by biological replicate and the Seurat function FindMarkers was used to identify differentially expressed genes in Mac2 and Mac3 cells using DESeq2.

In the Jerby-Anon et al. dataset, the macrophage compartment was subsetted and used for downstream analysis. The bottom 5% of expressed genes were filtered out before calculating the mean variance and expression. A bin size of 250 windows was used for calculating running mean variance. The ADCP gene module was mapped to human orthologs before calculating the ADCP gene module score using the Seurat package. Significance between the distance from smoothed variance between ADCP genes and all genes was determined using a Wilcoxon rank-sum test.

As a control for the Pearson correlation calculation between ADCP gene module score and PC2, 1000 sets of control genes were randomly selected from bins of similar expression to ADCP genes (genes were split into 25 bins based on mean expression quantiles). For each set of control genes, the gene module score was calculated using the Seurat package, and the Pearson correlation calculation between each control gene set and PC2 was calculated. This same analysis was performed on the glioblastoma and synovial sarcoma datasets.

## REFERENCES

1. A-Gonzalez, N., et al. Phagocytosis imprints heterogeneity in tissue-resident macrophages. J. Exp. Med. 214, 1281–1296 (2017).

2. Hirayama, D., Iida, T. & Nakase, H. The phagocytic function of macrophage-enforcing innate immunity and tissue homeostasis. Int. J. Mol. Sci. 19, 92 (2017).

3. Aderem, A. & Underhill, D. M. Mechanisms of phagocytosis in macrophages. Annu. Rev. Immunol. 17, 593–623 (1999).

4. Canetti, C., Hu, B., Curtis, J. L. & Peters-Golden, M. Syk activation is a leukotriene B4-regulated event involved in macrophage phagocytosis of IgG-coated targets but not apoptotic cells. Blood 102, 1877–1883 (2003).

5. Sánchez-Mejorada, G. & Rosales, C. Fcgamma receptor-mediated mitogen-activated protein kinase activation in monocytes is independent of Ras. J. Biol. Chem. 273, 27610– 27619 (1998).

6. Bournazos, S., Gupta, A. & Ravetch, J. V. The role of IgG Fc receptors in antibody-dependent enhancement. Nat. Rev. Immunol. 20, 633–643 (2020).

7. Kiefer, F. et al. The Syk protein tyrosine kinase is essential for Fcgamma receptor signaling in macrophages and neutrophils. Mol. Cell. Biol. 18, 4209–4220 (1998).

8. Ellis, S., Lin, E. J. & Tartar, D. Immunology of wound healing. Curr. Dermatol. Rep. 7, 350–358 (2018).

9. Liebold, I. et al. Apoptotic cell identity induces distinct functional responses to IL-4 in efferocytic macrophages. Science 384, eabo7027 (2024).

10. Gerlach, B. D. et al. Efferocytosis induces macrophage proliferation to help resolve tissue injury. Cell Metab. 33, 2445–2463.e8 (2021).

11. Yin, C. & Heit, B. Cellular responses to the efferocytosis of apoptotic cells. Front. Immunol. 12, 631714 (2021).

12. Fadok, V. A. et al. Macrophages that have ingested apoptotic cells in vitro inhibit proinflammatory cytokine production through autocrine/paracrine mechanisms involving TGF-beta, PGE2, and PAF. J. Clin. Invest. 101, 890–898 (1998).

13. Yagi, T. et al. Tumour-associated macrophages are associated with poor prognosis and programmed death ligand 1 expression in oesophageal cancer. Eur. J. Cancer 111, 38–49 (2019).

14. Zhang, J. et al. High infiltration of tumor-associated macrophages influences poor prognosis in human gastric cancer patients, associates with the phenomenon of EMT. Medicine (Baltimore) 95, e2636 (2016).

15. Gwak, J. M., Jang, M. H., Kim, D. I., Seo, A. N. & Park, S. Y. Prognostic value of tumor-associated macrophages according to histologic locations and hormone receptor status in breast cancer. PLoS One 10, e0125728 (2015).

16. Yuan, Z.-Y., Luo, R.-Z., Peng, R.-J., Wang, S.-S. & Xue, C. High infiltration of tumor-associated macrophages in triple-negative breast cancer is associated with a higher risk of distant metastasis. Onco. Targets. Ther. 7, 1475–1480 (2014).

17. Neophytou, C. M. et al. The role of tumor-associated myeloid cells in modulating cancer therapy. Front. Oncol. 10, 899 (2020).

18. Noy, R. & Pollard, J. W. Tumor-associated macrophages: from mechanisms to therapy. Immunity 41, 49–61 (2014).

19. Pittet, M. J., Michielin, O. & Migliorini, D. Clinical relevance of tumour-associated macrophages. Nat. Rev. Clin. Oncol. 19, 402–421 (2022).

20. Bied, M., Ho, W. W., Ginhoux, F. & Blériot, C. Roles of macrophages in tumor development: a spatiotemporal perspective. Cell. Mol. Immunol. 20, 983–992 (2023).

21. Pan, Y., Yu, Y., Wang, X. & Zhang, T. Tumor-associated macrophages in tumor immunity. Front. Immunol. 11, 583084 (2020).

22. Park, M. D. et al. TREM2 macrophages drive NK cell paucity and dysfunction in lung cancer. Nat. Immunol. 24, 792–801 (2023).

23. Gonzalez, M. A. et al. Phagocytosis increases an oxidative metabolic and immune suppressive signature in tumor macrophages. J. Exp. Med. 220, (2023).

24. Astuti, Y. et al. Efferocytosis reprograms the tumor microenvironment to promote pancreatic cancer liver metastasis. *Nat*. Cancer 5, 774–790 (2024).

25. Molgora, M. et al. TREM2 modulation remodels the tumor myeloid landscape enhancing anti-PD-1 immunotherapy. Cell 182, 886–900.e17 (2020).

26. Azizi, E. et al. Single-cell map of diverse immune phenotypes in the breast tumor microenvironment. Cell 174, 1293–1308.e36 (2018).

27. Zilionis, R. et al. Single-cell transcriptomics of human and mouse lung cancers reveals conserved myeloid populations across individuals and species. Immunity 50, 1317–1334.e10 (2019).

28. Casanova-Acebes, M. et al. Tissue-resident macrophages provide a pro-tumorigenic niche to early NSCLC cells. Nature 595, 578–584 (2021).

29. Mulder, K. et al. Cross-tissue single-cell landscape of human monocytes and macrophages in health and disease. Immunity 54, 1883–1900.e5 (2021).

30. Cheng, S. et al. A pan-cancer single-cell transcriptional atlas of tumor infiltrating myeloid cells. Cell 184, 792–809.e23 (2021).

31. Ma, R.-Y., Black, A. & Qian, B.-Z. Macrophage diversity in cancer revisited in the era of single-cell omics. Trends Immunol. 43, 546–563 (2022).

32. Gholamin, S. et al. Disrupting the CD47-SIRPα anti-phagocytic axis by a humanized anti-CD47 antibody is an efficacious treatment for malignant pediatric brain tumors. Sci. Transl. Med. 9, (2017).

33. Chao, M. P. et al. Anti-CD47 antibody synergizes with rituximab to promote phagocytosis and eradicate non-Hodgkin lymphoma. Cell 142, 699–713 (2010).

34. Scott, A. M., Wolchok, J. D. & Old, L. J. Antibody therapy of cancer. Nat. Rev. Cancer 12, 278–287 (2012).

35. Sliwkowski, M. X. & Mellman, I. Antibody therapeutics in cancer. Science 341, 1192–1198 (2013).

36. Glennie, M. J., French, R. R., Cragg, M. S. & Taylor, R. P. Mechanisms of killing by anti-CD20 monoclonal antibodies. Mol. Immunol. 44, 3823–3837 (2007).

37. Oflazoglu, E. et al. Macrophages and Fc-receptor interactions contribute to the antitumour activities of the anti-CD40 antibody SGN-40. Br. J. Cancer 100, 113–117 (2009).

38. Su, S. et al. Immune checkpoint inhibition overcomes ADCP-induced immunosuppression by macrophages. Cell 175, 442–457.e23 (2018).

39. Kamber, R. A. et al. Inter-cellular CRISPR screens reveal regulators of cancer cell phagocytosis. Nature 597, 549–554 (2021).

40. Gray, M. A. et al. Phagocytosis enhances lysosomal and bactericidal properties by activating the transcription factor TFEB. Curr. Biol. 26, 1955–1964 (2016).

41. Tseng, D. et al. Anti-CD47 antibody-mediated phagocytosis of cancer by macrophages primes an effective antitumor T-cell response. Proc. Natl. Acad. Sci. U. S. A. 110, 11103–11108 (2013).

42. Muliaditan, T. et al. Macrophages are exploited from an innate wound healing response to facilitate cancer metastasis. Nat. Commun. 9, 2951 (2018).

43. Patsalos, A. et al. The BACH1-HMOX1 regulatory axis is indispensable for proper macrophage subtype specification and skeletal muscle regeneration. J. Immunol. 203, 1532– 1547 (2019).

44. Grochot-Przeczek, A. et al. Heme oxygenase-1 accelerates cutaneous wound healing in mice. PLoS One 4, e5803 (2009).

45. Molgora, M., Liu, Y. A., Colonna, M. & Cella, M. TREM2: A new player in the tumor microenvironment. Semin. Immunol. 67, 101739 (2023).

46. Katzenelenbogen, Y. et al. Coupled scRNA-seq and intracellular protein activity reveal an immunosuppressive role of TREM2 in cancer. Cell 182, 872–885.e19 (2020).

47. Pesce, J. T., et al. Arginase-1-expressing macrophages suppress Th2 cytokine-driven inflammation and fibrosis. PLoS Pathog. 5, e1000371 (2009).

48. Cai, W. et al. STAT6/Arg1 promotes microglia/macrophage efferocytosis and inflammation resolution in stroke mice. JCI Insight 4, (2019).

49. Bill, R. et al. CXCL9:SPP1 macrophage polarity identifies a network of cellular programs that control human cancers. Science 381, 515–524 (2023).

50. Matsubara, E. et al. SPP1 derived from macrophages is associated with a worse clinical course and chemo-resistance in lung adenocarcinoma. Cancers (Basel) 14, 4374 (2022).

51. Savina, A. et al. The small GTPase Rac2 controls phagosomal alkalinization and antigen crosspresentation selectively in CD8(+) dendritic cells. Immunity 30, 544–555 (2009).

52. Hoppe, A. D. & Swanson, J. A. Cdc42, Rac1, and Rac2 display distinct patterns of activation during phagocytosis. Mol. Biol. Cell 15, 3509–3519 (2004).

53. Mao, Y. & Finnemann, S. C. Regulation of phagocytosis by Rho GTPases. Small GTPases 6, 89–99 (2015).

54. Zhang, S. et al. Efferocytosis fuels requirements of fatty acid oxidation and the electron transport chain to polarize macrophages for tissue repair. Cell Metab. 29, 443–456.e5 (2019).

55. Jin, Z., Wei, W., Yang, M., Du, Y. & Wan, Y. Mitochondrial complex I activity suppresses inflammation and enhances bone resorption by shifting macrophage-osteoclast polarization. Cell Metab. 20, 483–498 (2014).

56. Viola, A., Munari, F., Sánchez-Rodríguez, R., Scolaro, T. & Castegna, A. The metabolic signature of macrophage responses. Front. Immunol. 10, 1462 (2019).

57. Daniel, B. et al. Macrophage inflammatory and regenerative response periodicity is programmed by cell cycle and chromatin state. Mol. Cell 83, 121–138.e7 (2023).

58. Lavin, Y. et al. Tissue-resident macrophage enhancer landscapes are shaped by the local microenvironment. Cell 159, 1312–1326 (2014).

59. Gosselin, D. et al. Environment drives selection and function of enhancers controlling tissue-specific macrophage identities. Cell 159, 1327–1340 (2014).

60. Alam, J. et al. Nrf2, a Cap’n’Collar transcription factor, regulates induction of the heme oxygenase-1 gene. J. Biol. Chem. 274, 26071–26078 (1999).

61. Wang, P. et al. Macrophage achieves self-protection against oxidative stress-induced ageing through the Mst-Nrf2 axis. Nat. Commun. 10, 755 (2019).

62. Virág, L., Jaén, R. I., Regdon, Z., Boscá, L. & Prieto, P. Self-defense of macrophages against oxidative injury: Fighting for their own survival. Redox Biol. 26, 101261 (2019).

63. Kadl, A. et al. Identification of a novel macrophage phenotype that develops in response to atherogenic phospholipids via Nrf2. Circ. Res. 107, 737–746 (2010).

64. Hegde, S. et al. Myeloid progenitor dysregulation fuels immunosuppressive macrophages in tumors. *bioRxivorg* 2024.06.24.600383 (2024).

65. Beury, D. W. et al. Myeloid-derived suppressor cell survival and function are regulated by the transcription factor Nrf2. J. Immunol. 196, 3470–3478 (2016).

66. Piantadosi, C. A. et al. Heme oxygenase-1 couples activation of mitochondrial biogenesis to anti-inflammatory cytokine expression. J. Biol. Chem. 286, 16374–16385 (2011).

67. Reichard, J. F., Motz, G. T. & Puga, A. Heme oxygenase-1 induction by NRF2 requires inactivation of the transcriptional repressor BACH1. Nucleic Acids Res. 35, 7074–7086 (2007).

68. Loboda, A., Damulewicz, M., Pyza, E., Jozkowicz, A. & Dulak, J. Role of Nrf2/HO-1 system in development, oxidative stress response and diseases: an evolutionarily conserved mechanism. Cell. Mol. Life Sci. 73, 3221–3247 (2016).

69. Palmieri, M. et al. Characterization of the CLEAR network reveals an integrated control of cellular clearance pathways. Hum. Mol. Genet. 20, 3852–3866 (2011).

70. Piccolo, V. et al. Opposing macrophage polarization programs show extensive epigenomic and transcriptional cross-talk. Nat. Immunol. 18, 530–540 (2017).

71. Krause, C. D., He, W., Kotenko, S. & Pestka, S. Modulation of the activation of Stat1 by the interferon-gamma receptor complex. Cell Res. 16, 113–123 (2006).

72. Udofa, E. A. et al. The transcription factor C/EBP-β mediates constitutive and LPS-inducible transcription of murine SerpinB2. PLoS One 8, e57855 (2013).

73. Liu, Y.-W., Chen, C.-C., Tseng, H.-P. & Chang, W.-C. Lipopolysaccharide-induced transcriptional activation of interleukin-10 is mediated by MAPK- and NF-kappaB-induced CCAAT/enhancer-binding protein delta in mouse macrophages. Cell. Signal. 18, 1492– 1500 (2006).

74. Alam, T., An, M. R. & Papaconstantinou, J. Differential expression of three C/EBP isoforms in multiple tissues during the acute phase response. J. Biol. Chem. 267, 5021–5024 (1992).

75. Mishra, R. K., Potteti, H. R., Tamatam, C. R., Elangovan, I. & Reddy, S. P. C-Jun is required for nuclear factor-κB-dependent, LPS-stimulated Fos-related antigen-1 transcription in alveolar macrophages. Am. J. Respir. Cell Mol. Biol. 55, 667–674 (2016).

76. Puertollano, R., Ferguson, S. M., Brugarolas, J. & Ballabio, A. The complex relationship between TFEB transcription factor phosphorylation and subcellular localization. EMBO J. 37, (2018).

77. Carey, K. L. et al. TFEB transcriptional responses reveal negative feedback by BHLHE40 and BHLHE41. Cell Rep. 33, 108371 (2020).

78. Pastore, N. et al. TFEB and TFE3 cooperate in the regulation of the innate immune response in activated macrophages. Autophagy 12, 1240–1258 (2016).

79. Nair, S. et al. Transcription factor stoichiometry, motif affinity and syntax regulate single-cell chromatin dynamics during fibroblast reprogramming to pluripotency. bioRxiv (2023) doi:10.1101/2023.10.04.560808.

80. Pampari, A. et al. ChromBPNet: bias factorized, base-resolution deep learning models of chromatin accessibility reveal cis-regulatory sequence syntax, transcription factor footprints and regulatory variants. bioRxivorg (2025) doi:10.1101/2024.12.25.630221.

81. Durdu, S. et al. Chromatin-dependent motif syntax defines differentiation trajectories. *bioRxiv* 2024.08.05.606702 (2024) doi:10.1101/2024.08.05.606702.

82. Kim, D. S. et al. The dynamic, combinatorial cis-regulatory lexicon of epidermal differentiation. Nat. Genet. 53, 1564–1576 (2021).

83. Shrikumar, A. et al. Technical note on Transcription Factor Motif Discovery from Importance Scores (TF-MoDISco) version 0.5.6.5. *arXiv [cs.LG]* (2018).

84. Liguori, M. et al. The soluble glycoprotein NMB (GPNMB) produced by macrophages induces cancer stemness and metastasis via CD44 and IL-33. Cell. Mol. Immunol. 18, 711– 722 (2021).

85. Zhou, L. et al. Glycoprotein non-metastatic melanoma protein b (Gpnmb) is highly expressed in macrophages of acute injured kidney and promotes M2 macrophages polarization. Cell. Immunol. 316, 53–60 (2017).

86. Saade, M., Araujo de Souza, G., Scavone, C. & Kinoshita, P. F. The role of GPNMB in inflammation. Front. Immunol. 12, 674739 (2021).

87. Ripoll, V. M. et al. Microphthalmia transcription factor regulates the expression of the novel osteoclast factor GPNMB. Gene 413, 32–41 (2008).

88. Hong, S.-B. et al. Inactivation of the FLCN tumor suppressor gene induces TFE3 transcriptional activity by increasing its nuclear localization. PLoS One 5, e15793 (2010).

89. Vorselen, D. et al. Cell surface receptors TREM2, CD14 and integrin α_M_β_2_ drive sinking engulfment in phosphatidylserine-mediated phagocytosis. *bioRxiv* 2022.07.30.502145 (2022) doi:10.1101/2022.07.30.502145.

90. Underhill, D. M. et al. The Toll-like receptor 2 is recruited to macrophage phagosomes and discriminates between pathogens. Nature 401, 811–815 (1999).

91. Yurdagul, A., Jr et al. Macrophage metabolism of apoptotic cell-derived arginine promotes continual efferocytosis and resolution of injury. Cell Metab. 31, 518–533.e10 (2020).

92. Ngai, D., Schilperoort, M. & Tabas, I. Efferocytosis-induced lactate enables the proliferation of pro-resolving macrophages to mediate tissue repair. Nat. Metab. 5, 2206– 2219 (2023).

93. Boada-Romero, E., Martinez, J., Heckmann, B. L. & Green, D. R. The clearance of dead cells by efferocytosis. Nat. Rev. Mol. Cell Biol. 21, 398–414 (2020).

94. Tufan, T. et al. Rapid unleashing of macrophage efferocytic capacity via transcriptional pause release. Nature 628, 408–415 (2024).

95. Sumitomo, R. et al. M2-like tumor-associated macrophages promote epithelial-mesenchymal transition through the transforming growth factor β/Smad/zinc finger e-box binding homeobox pathway with increased metastatic potential and tumor cell proliferation in lung squamous cell carcinoma. Cancer Sci. 114, 4521–4534 (2023).

96. Chen, X. et al. Tumor-associated macrophages promote epithelial-mesenchymal transition and the cancer stem cell properties in triple-negative breast cancer through CCL2/AKT/β-catenin signaling. Cell Commun. Signal. 20, 92 (2022).

97. Han, K. et al. CRISPR screens in cancer spheroids identify 3D growth-specific vulnerabilities. Nature 580, 136–141 (2020).

98. Kumar, V., Patel, S., Tcyganov, E. & Gabrilovich, D. I. The nature of myeloid-derived suppressor cells in the tumor microenvironment. Trends Immunol. 37, 208–220 (2016).

99. Kohli, K., Pillarisetty, V. G. & Kim, T. S. Key chemokines direct migration of immune cells in solid tumors. Cancer Gene Ther. 29, 10–21 (2022).

100. Remmerie, A. & Scott, C. L. Macrophages and lipid metabolism. Cell. Immunol. 330, 27– 42 (2018).

101. Wu, F. et al. Single-cell profiling of tumor heterogeneity and the microenvironment in advanced non-small cell lung cancer. Nat. Commun. 12, 2540 (2021).

102. Settembre, C. et al. TFEB links autophagy to lysosomal biogenesis. Science 332, 1429– 1433 (2011).

103. Sardiello, M. et al. A gene network regulating lysosomal biogenesis and function. Science 325, 473–477 (2009).

104. Roczniak-Ferguson, A. et al. The transcription factor TFEB links mTORC1 signaling to transcriptional control of lysosome homeostasis. Sci. Signal. 5, ra42 (2012).

105. Visvikis, O. et al. Innate host defense requires TFEB-mediated transcription of cytoprotective and antimicrobial genes. Immunity 40, 896–909 (2014).

106. van den Broek, T. J. M. et al. Single-cell spatial analysis of pediatric high-grade glioma reveals a novel population of SPP1^+^/GPNMB^+^myeloid cells with immunosuppressive and tumor-promoting capabilities. *bioRxiv* 2025.03.18.643953 (2025) doi:10.1101/2025.03.18.643953.

107. Masetti, M. et al. Lipid-loaded tumor-associated macrophages sustain tumor growth and invasiveness in prostate cancer. J. Exp. Med. 219, (2022).

108. Jerby-Arnon, L. et al. A cancer cell program promotes T cell exclusion and resistance to checkpoint blockade. Cell 175, 984–997.e24 (2018).

109. Neftel, C. et al. An integrative model of cellular states, plasticity, and genetics for glioblastoma. Cell 178, 835–849.e21 (2019).

110. Jerby-Arnon, L. et al. Opposing immune and genetic mechanisms shape oncogenic programs in synovial sarcoma. Nat. Med. 27, 289–300 (2021).

111. Winslow, M. M. et al. Suppression of lung adenocarcinoma progression by Nkx2-1. Nature 473, 101–104 (2011).

112. Callari, M. et al. Computational approach to discriminate human and mouse sequences in patient-derived tumour xenografts. BMC Genomics 19, 19 (2018).

113. Dobin, A. et al. STAR: ultrafast universal RNA-seq aligner. Bioinformatics 29, 15–21 (2013).

114. Putri, G. H., Anders, S., Pyl, P. T., Pimanda, J. E. & Zanini, F. Analysing high-throughput sequencing data in Python with HTSeq 2.0. Bioinformatics 38, 2943–2945 (2022).

115. Love, M. I., Huber, W. & Anders, S. Moderated estimation of fold change and dispersion for RNA-seq data with DESeq2. Genome Biol. 15, 550 (2014).

116. Pinney, J. J. et al. Macrophage hypophagia as a mechanism of innate immune exhaustion in mAb-induced cell clearance. Blood 136, 2065–2079 (2020).

117. Van Dyken, S. J. & Locksley, R. M. Interleukin-4- and interleukin-13-mediated alternatively activated macrophages: roles in homeostasis and disease. Annu. Rev. Immunol. 31, 317–343 (2013).

118. Gordon, S. & Martinez, F. O. Alternative activation of macrophages: mechanism and functions. Immunity 32, 593–604 (2010).

119. Bosurgi, L. et al. Macrophage function in tissue repair and remodeling requires IL-4 or IL-13 with apoptotic cells. Science 356, 1072–1076 (2017).

120. Loi, S. et al. Tumor infiltrating lymphocytes are prognostic in triple negative breast cancer and predictive for trastuzumab benefit in early breast cancer: results from the FinHER trial. Ann. Oncol. 25, 1544–1550 (2014).

121. Park, S. et al. The therapeutic effect of anti-HER2/neu antibody depends on both innate and adaptive immunity. Cancer Cell 18, 160–170 (2010).

122. Xu, M. et al. Intratumoral delivery of IL-21 overcomes anti-Her2/Neu resistance through shifting tumor-associated macrophages from M2 to M1 phenotype. J. Immunol. 194, 4997– 5006 (2015).

123. Voll, R. E. et al. Immunosuppressive effects of apoptotic cells. Nature 390, 350–351 (1997).

124. Toossi, Z. et al. Induction of serine protease inhibitor 9 by Mycobacterium tuberculosis inhibits apoptosis and promotes survival of infected macrophages. J. Infect. Dis. 205, 144– 151 (2012).

